# Mutation and cell state compatibility is required and targetable in Ph*+* acute lymphoblastic leukemia minimal residual disease

**DOI:** 10.1101/2024.06.06.597767

**Authors:** Peter S. Winter, Michelle L. Ramseier, Andrew W. Navia, Sachit Saksena, Haley Strouf, Nezha Senhaji, Alan DenAdel, Mahnoor Mirza, Hyun Hwan An, Laura Bilal, Peter Dennis, Catharine S. Leahy, Kay Shigemori, Jennyfer Galves-Reyes, Ye Zhang, Foster Powers, Nolawit Mulugeta, Alejandro J. Gupta, Nicholas Calistri, Alex Van Scoyk, Kristen Jones, Huiyun Liu, Kristen E. Stevenson, Siyang Ren, Marlise R. Luskin, Charles P. Couturier, Ava P. Amini, Srivatsan Raghavan, Robert J. Kimmerling, Mark M. Stevens, Lorin Crawford, David M. Weinstock, Scott R. Manalis, Alex K. Shalek, Mark A. Murakami

## Abstract

Efforts to cure BCR::ABL1 B cell acute lymphoblastic leukemia (Ph+ ALL) solely through inhibition of ABL1 kinase activity have thus far been insufficient despite the availability of tyrosine kinase inhibitors (TKIs) with broad activity against resistance mutants. The mechanisms that drive persistence within minimal residual disease (MRD) remain poorly understood and therefore untargeted. Utilizing 13 patient-derived xenograft (PDX) models and clinical trial specimens of Ph+ ALL, we examined how genetic and transcriptional features co-evolve to drive progression during prolonged TKI response. Our work reveals a landscape of cooperative mutational and transcriptional escape mechanisms that differ from those causing resistance to first generation TKIs. By analyzing MRD during remission, we show that the same resistance mutation can either increase or decrease cellular fitness depending on transcriptional state. We further demonstrate that directly targeting transcriptional state-associated vulnerabilities at MRD can overcome BCR::ABL1 independence, suggesting a new paradigm for rationally eradicating MRD prior to relapse. Finally, we illustrate how cell mass measurements of leukemia cells can be used to rapidly monitor dominant transcriptional features of Ph+ ALL to help rationally guide therapeutic selection from low-input samples.

**HIGHLIGHTS:**

- Relapse after remission on TKI can harbor mutations in ABL1, RAS, or neither
- Mutations and development-like cell state dictate fitness in residual disease
- Co-targeting cell state and ABL1 markedly reduces MRD
- Biophysical measurements provide an integrative, rapid measurement of cell state

## INTRODUCTION

A large fraction of patients with cancer achieve complete remission at some point during their course of therapy, either through surgery, chemotherapy, radiation, or a combination thereof. Nevertheless, many of these patients relapse or progress owing to a small pool of remaining cancer cells commonly referred to as minimal residual disease (MRD). This is even true for cancers with clear, targetable oncogene dependencies such as BCR::ABL1-rearranged B cell acute lymphoblastic leukemia (Ph+ ALL). Despite highly effective tyrosine kinase inhibitors (TKI) with potent activity against multiple resistance-conferring point mutations in BCR::ABL1, relapse during single-agent treatment is nearly universal.^1,2,3^ Unfortunately, accumulating evidence casts doubt on the potential for up-front combinations of next-generation TKIs to fully overcome subclonal heterogeneity and thereby eradicate MRD.^4^

While most patients with BCR::ABL1-driven disease relapse with kinase domain mutations, 30-40% of patients progress with BCR::ABL1-independent mechanisms that are poorly understood.^5^ Previous studies have identified developmental heterogeneity across ALL,^6,7^ as well as in Ph+ ALL specifically.^8,9^ This developmental heterogeneity has also been linked to treatment response for multiple classes of inhibitors.^6,7,10^ Recent work specifically in Ph+ ALL examined developmental subtypes that align with earlier (Early-Pro) and later developmental (Late-Pro) B cell features, finding that the former was associated with poor overall survival upon treatment with the first-generation TKI imatinib.^8^ Commitment to earlier or later stages of development has been associated with cooperating alterations in lineage-defining transcription factors (*EBF1* deletion or deletions in *IKZF1, PAX5,* and *CDKN2A,* respectively), suggesting that developmental state adherence – and its associated therapeutic response – may be mutationally driven and static upon leukemic transformation.^8,9^ However, other studies have nominated the potential for a leukemia’s dominant developmental states to shift in response to therapeutic pressure. Illustratively, non-mutational mechanisms of chemotherapy resistance have been observed in ALL patient-derived xenografts (PDXs), whereby leukemia cells transiently adopt a dormant, stem-like state at MRD;^11^ others have demonstrated post-treatment shifts in the abundance of dormant subpopulations mimicking earlier developmental stages.^7^ It has also been suggested that TKI-resistant Ph+ ALL cells in a later developmental state proliferate by activating signaling that typically occurs downstream of the pre-B cell receptor (pre-BCR), despite the absence of a functionally expressed pre-BCR in Ph+ ALL.^10,12^ It remains unclear which attributes allow for persistence during remission and if mutational or developmental phenotypes are the dominant drivers of resistance.

Accordingly, resistance to ABL1 TKIs is multifactorial and extends beyond ABL1 resistance mutations, suggesting that informed strategies to convert deep remissions into cures may require incorporating orthogonal measurements of the non-genetic determinants of cellular state (e.g. via single-cell transcriptomics).^13,14,15^ However, there are limited studies describing how mutations participate (or clash) with these additional cellular features to drive persistence and clonal expansion under TKI pressure. Though recent evidence from our group and others indicates that some mutations are enriched in specific transcriptional backgrounds,^10,16,17,18,19^ the relative importance of mutational and transcriptional drivers to MRD persistence and relapse is not known. Furthermore, there are significant technical challenges associated with isolating and profiling rare residual cells that have limited their characterization largely to mutational profiling – a problem affecting essentially all cancer types.^13,20,21^ While MRD enumeration and mutational monitoring have been used to some clinical benefit,^22,23,24^ the translational utility of understanding non-mutational attributes from these rare cells has yet to be demonstrated.^21^ These constraints, coupled with the heterogeneity among MRD phenotypes both within and between patients, have historically made it difficult to nominate specific therapeutic strategies to combat MRD. We and others previously proposed that direct interrogation of MRD cells to identify dependencies for individual patients could offer clinical benefit if approaches existed to define those dependences in “real-time”.^21,25^ This would require a rapid strategy applicable to individual cells that could distinguish patients most likely to respond to one of several available therapeutic options.

Here, to better understand how both mutational and transcriptional variation coordinate to drive relapse within MRD, we defined the biology of Ph+ ALL cells at different stages of treatment and across a diversity of models and human patients. We reveal unique and targetable characteristics of Ph+ ALL MRD and nominate combination strategies to eradicate residual disease.

## RESULTS

### Modeling disease kinetics in response to combination TKI in Ph+ ALL PDX models

Although treatment with allosteric BCR::ABL1 inhibitors drives deep remissions in patients, nearly all will relapse if not consolidated with allogeneic stem cell transplantation. The recent development of asciminib (ABL001), an allosteric inhibitor of BCR::ABL1,^26^ created the first opportunity to address whether dual inhibition of BCR::ABL1 could eradicate Ph+ leukemias (**Figure 1A**). We combined orthosteric (ponatinib; 40 mg/kg/day) and allosteric (asciminib; 30 mg/kg/day) inhibitors in a diverse cohort of Ph+ ALL PDX models (n=13 models;^27^ 190 mice total) to assess how pre-existing clinical and molecular features would dictate response to sustained oncogene withdrawal within a statistically powered, phase II-like preclinical trial (**Figures 1A & 1B;** see **Methods** and **Tables S1 & S2**). All mice receiving ponatinib or combination therapy, and 92% of subjects receiving asciminib monotherapy who survived beyond one week achieved complete remission (CR), corroborating the dependence of these leukemias on BCR::ABL1 (**Figures S1A & S1B**). The durations of remission with ponatinib-based regimens exceeded those of asciminib monotherapy, but we observed no difference between the combination and ponatinib monotherapy arms (p=0.70; **Figures 1C & S1C**). Notably, survival outcomes between mice on each treatment arm did not correlate with PDX line characteristics associated with inferior treatment response in other contexts, such as increased prior lines of therapy,^28^ *IKZF1* deletion,^29^ and pre-existing *ABL1* resistance mutations (**Figure S1D**).^2,30^ All mice were ultimately euthanized, either for disease progression or clinical toxicity. Even the 7 mice euthanized for clinical toxicity after achieving a durable response – three of whom maintained CR for >12 months on study (**Figure S1B**) – harbored residual ALL in the bone marrow and/or spleen when sacrificed. These data demonstrate that while single-agent ponatinib and combination therapy confer deep and prolonged clinical remissions, BCR::ABL1 inhibition alone was insufficient to fully eradicate human leukemias *in vivo*.

**Figure 1.**
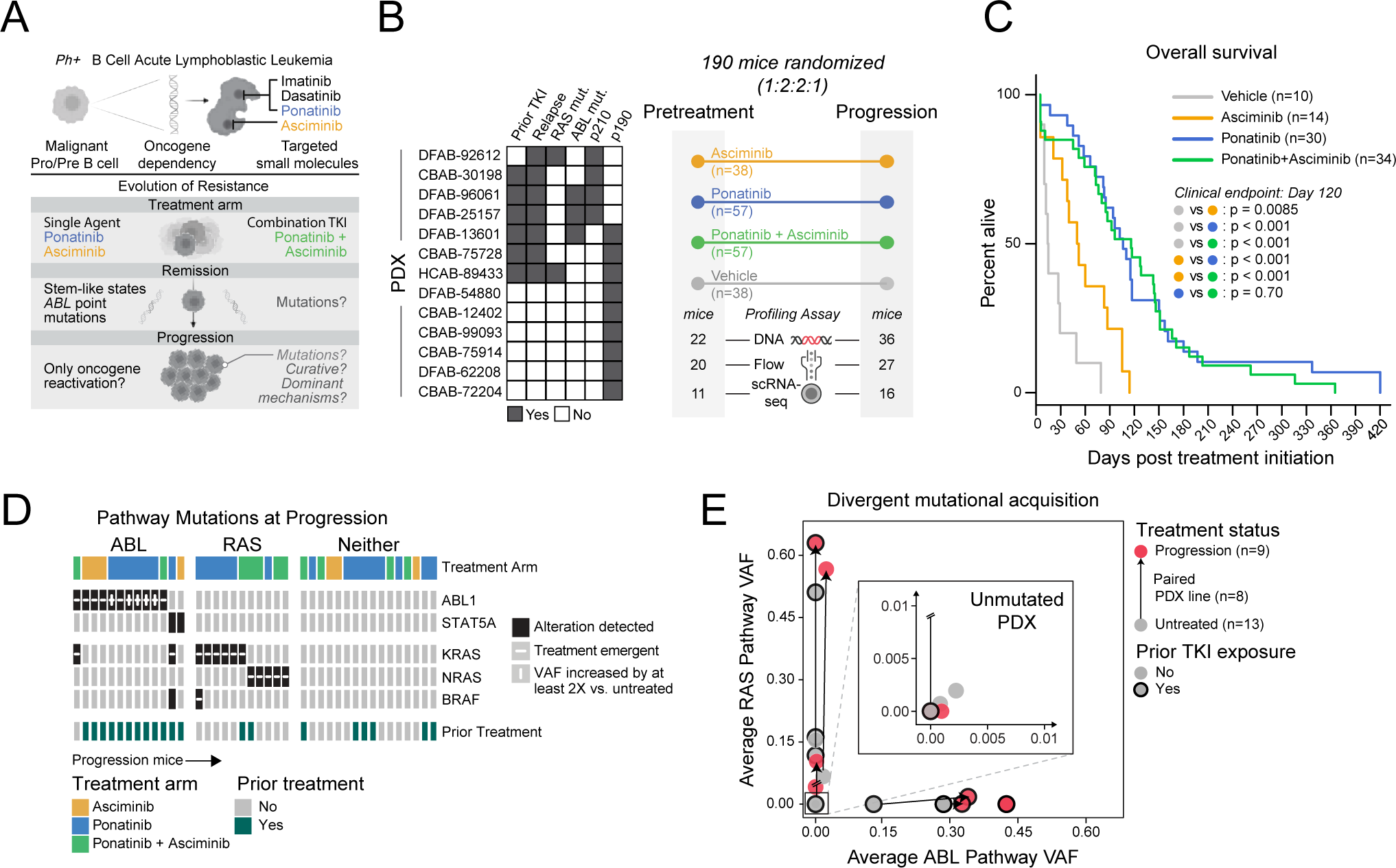
Genetic mechanisms of resistance to oncogene inhibition in Ph+ ALL. **(A)** Motivation for evaluating efficacy and mechanisms of resistance to combination TKI therapy in Ph+ ALL. **(B)** Patient characteristics of the 13 PDX models used in the study and Phase II-like randomized *in vivo* trial design. Number of mice examined by genetic profiling, immunophenotyping (“Flow”), or scRNA-seq at pre-treatment and progression time points. For characteristics of patients from whom PDX lines were derived (Table S2): “TKI”=prior patient exposure to tyrosine kinase inhibitor; “relapse”=patient tumor at progression; “mut”=mutant (non-*BCR::ABL1*); “p210” and “p190”=p210 and p190 *BCR::ABL1* isoforms, respectively. **(C)** Overall survival across treatment arms in Phase II-like study; p-values from Cox regression analysis at clinical end-point (day 120) are indicated for each pairwise comparison between treatment arms. **(D)** ABL and RAS pathway detected alterations in Phase II-like study tumors at progression (n=40). Treatment emergent mutations indicated when mice from the same PDX line were profiled at pretreatment (see full alteration details in Figure S2A and Table S3). Prior treatment indicates mice whose PDX lines were derived from patients with prior TKI and chemotherapy exposure. “VAF”=variant allele frequency. **(E)** Average VAF for mutations along RAS (y-axis) or ABL (x-axis) pathways, averaged across mice in each PDX line at pretreatment or progression. Arrows link pretreatment and progression average VAFs from the same PDX line. PDX lines derived from patients with prior TKI exposure are outlined in black. Inset highlights a subset of PDX model timepoints where no (n=4 pretreatment, n=1 progression) or few mutations were detected in either pathway. *See also Figures S1 & S2; Tables S1, S2 & S3*.

### Divergent mutational patterns upon oncogene inhibition in Ph+ B-ALL

To chart landscapes of genetic resistance to single agent and combination TKI in Ph+ ALL, we sequenced 142 PDX samples (74 trial and 68 other TKI-treated leukemias) and examined patterns of acquisition within known driver mutations in ALL across multiple phases of treatment (**Figure S2A**; **Table S3;** see **Methods**). In general, alterations in ABL1 or RAS pathway genes consistently emerged upon therapeutic pressure compared to mutations affecting B cell survival, lineage commitment, or cell cycle control (**Figures S2A & S2B**). Of relapsed leukemias, 35% harbored mutations in BCR::ABL1, frequently compound mutations involving T315I plus at least one other high-level resistance mutation (e.g., Y253H, F311L, F359V) or an activating mutation in *STAT5A* (collectively termed ‘ABL pathway’ mutations). A separate 24% relapsed with activating mutations in RAS pathway genes – specifically *KRAS*, *NRAS*, *BRAF*, and/or *PTPN11* – representing emergent alternate pathway utilization in these oncogene-addicted leukemias (**Figures 1D & S2C**). Acquisition of driver pathway mutations was influenced by treatment arm – mice treated with asciminib predominantly acquired ABL pathway mutations at relapse, mice treated with ponatinib predominantly acquired RAS pathway mutations, and mice on the dual-treatment arm acquired mutations on either ABL or RAS pathways (**Figure S2B**). Samples harboring RAS pathway mutations were mutually exclusive with those involving ABL pathway mutations within each PDX line at both pretreatment and progression time points (**Figure 1E**). The remaining tumors (41%) harbored no driver mutations in either ABL or RAS pathway genes, and the majority of these (74%) had no apparent genetic lesions explaining phenotypic resistance by whole exome sequencing (**Figures 1D, S2A, & S2C**). These data suggest three recurrent patterns for resistance whereby leukemias progress on therapy with either ABL pathway, RAS pathway, or no discernible gain-of-function mutations.

### Ph+ ALL leukemic cells are defined by hybrid developmental states

Given the lack of discernible mutation-driven resistance in a substantial fraction of our cohort (**Figures 1D & S2A**), we hypothesized that resistance to single-agent or combination TKI in Ph+ ALL may be understood best by characterizing both mutational and transcriptional state heterogeneity. To this end, we applied single-cell RNA-sequencing (scRNA-seq) to define transcriptional states in Ph+ ALL and identify leukemic phenotypes associated with progression. Using Seq-Well S^3,31^ we generated a dataset of 42,667 single-cell transcriptomes from 52 samples spanning 11 PDX lines from our phase II-like pre-clinical trial and 5 patients on a clinical trial testing dasatinib (a second-generation orthosteric BCR::ABL1 inhibitor) plus asciminib for previously untreated Ph+ ALL (**Figure 2A**; NCT02081378; see **Methods**). We then performed consensus non-negative matrix factorization (cNMF) over each leukemia in this dataset to identify intratumoral gene expression programs (GEPs; **Methods**). Hierarchical clustering of the 126 GEPs defined across individual leukemias revealed 7 shared patterns (meta-GEPs, or “mGEPs”) of covarying gene programming that were present in at least 8 samples (**Figures 2B, 2C & S3A; Table S4**). Two mGEPs were defined by genes associated with active stages of the cell cycle (e.g., *CENPF, MKI67, MCM6, E2F2*) and another mGEP specifically associated with MYC activity (e.g., *HSP90AB1, NME1*). The remaining four mGEPs associated with various stages of B cell development, either containing Pro-B cell genes (e.g., *DNTT*, *CSGALNACT1*), genes associated with later stages of B cell development — i.e., Pre-BII (e.g., *CD38, IRF4*), and Immature B (e.g., *CD79A, HLA-DPB1*) – or progenitor-associated genes co-expressed with Immature B genes (e.g., *CD44, CSF1R* and *HLA-DQA1, IRF8*). These data suggest that aspects of normal B cell development are captured as major axes of intratumoral transcriptional variation in Ph+ ALL.

**Figure 2.**
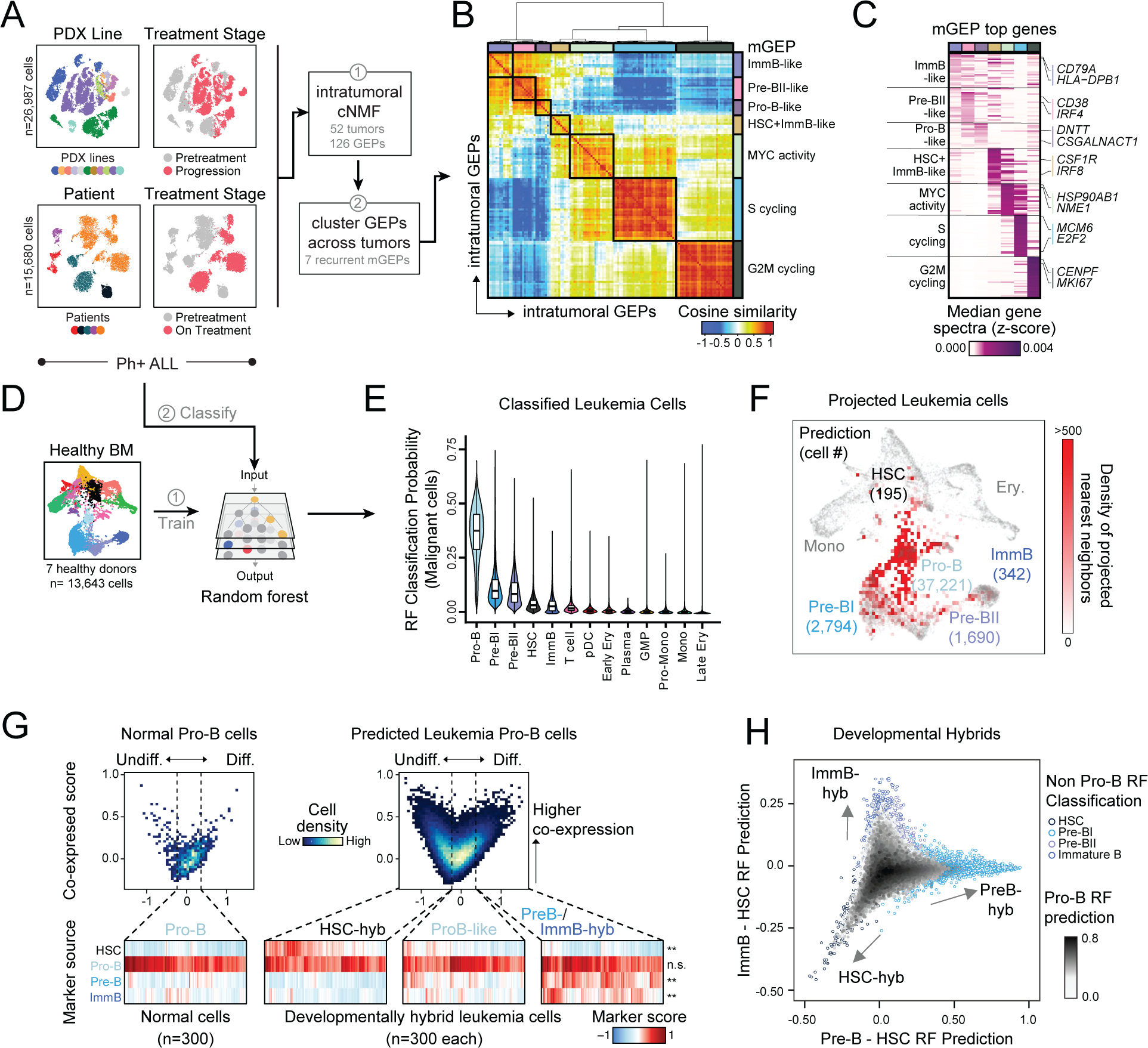
Hybrid developmental transcriptional states define B-ALL. **(A)** Overview of Ph+ ALL scRNA-seq data collected from PDX lines (n=26,987 cells across 11 PDX lines inclusive of 38 pretreatment and progression tumors) and patient biopsies (n=15,680 cells across 5 patients inclusive of 14 pretreatment and on-treatment tumors). **(B)** Unbiased factorization of leukemic scRNA-seq data with consensus non-negative matrix factorization (cNMF). Each row and column is an individual GEP and clustering is based on cosine similarity to find meta-programs (mGEPs; see Methods). “HSC”=hematopoietic stem cell; “ImmB”=Immature B. **(C)** Each mGEP annotated by the top 30 genes with the highest median cNMF gene spectra score across clustered intratumoral GEPs (Table S4). **(D)** Approach for supervised classification using a random forest (RF) classifier trained on healthy bone marrow (BM) scRNA-seq data. **(E)** Distribution (box plot and violin plot) of leukemia single-cell RF classification probabilities for each healthy BM cell type, ordered by median RF classification probability. “pDC”=plasmacytoid dendritic cell; “Ery”=erythroid; “Plasma”=plasma cell; “GMP”=granulocyte-monocyte progenitor; “Mono”=monocyte. **(F)** K-nearest neighbor (kNN) projection of all leukemia cells onto reference normal hierarchy, annotated by number of classified leukemic cells for each reference B cell lineage population. **(G)**Developmental marker gene co-expression in normal Pro-B cells (left) vs. leukemia cells classified as Pro-B cells (right). X-axis represents gene expression score difference between healthy HSC differentially expressed genes (undifferentiated) and the union of Pre-B and Immature B differentially expressed genes (more differentiated); y-axis represents each cell’s second highest healthy cell type marker expression score. 300 randomly-sampled single cells from each bin are shown below. P-values from ANOVA (\*\**p*<0.001) compare expression distribution in normal vs leukemic Pro-B cells for each normal cell type marker expression score (rows). **(H)** Leukemia cells plotted according to non Pro-B RF classification probabilities. Cells are colored by RF Pro-B classification probability (greyscale, fill) and cells are outlined by their classified cell type. *See also Figures S3-S6; Tables S4 & S5*.

In several cases, genes defining multiple B lineage developmental stages were enriched in the same GEP and co-expressed within individual leukemia cells (**Figures S3A & S3B**).^7,32^ We next sought to better understand these stage-specific “hybrid” expression patterns by utilizing a supervised machine learning approach to resolve the relationship between leukemia cells and nonmalignant B cell development. To enable this comparison, we first generated a reference dataset of human hematopoiesis from the bone marrow aspirates of healthy donors (n=7), profiling both sorted and unsorted fractions to ensure the proper B cell developmental populations were captured (**Figures S4A & S4B;** see **Methods**). By performing iterative clustering, we identified 13 cell types spanning the HSC progenitor, myeloid, erythroid, and lymphoid lineages (n = 13,643 cells; **Figures S4C & S4D**); each cell type population contained cells from at least 6 of 7 donors (**Figures S4E & S4F**). To enable leukemic cell reference mapping and comparison, we trained a random-forest (RF) classifier on the cell type-labeled reference scRNA-seq dataset using 10-fold cross-validation (**Figures 2D & S5A;** see **Methods**). We ensured this model was cueing on biologically-relevant expression patterns by using permutation tests to identify the top 200 features needed to accurately classify single-cell transcriptomes, as well as testing its accuracy on an external scRNA-seq dataset (**Figures S5B & S5C**).^16^ We then assigned individual B-ALL cells to their most likely developmental state using our RF classifier (**Figure 2D**). Across all malignant cells, the RF model assigned highest classification probabilities for the Pro-B cell type, followed by Pre-BI, Pre-BII, HSC, and Immature B cell types (**Figures 2E & 2F**); 1% of leukemia cells that classified into non-B lineage cell types, such as T cells, were poor quality and removed from downstream analyses (**Figure S5D**).

Corroborating our observations with NMF, marker genes that were restricted to individual stages of B cell development in healthy cells were routinely co-expressed in leukemia cells (**Figure 2G**). For example, within leukemic cells classified as Pro-B, we observed a dominant secondary RF classification probability for an earlier (HSC) or later (Pre-BI, Pre-BII, Immature B) stage of B cell development. We therefore characterized the transcriptional heterogeneity in Ph+ ALL as a continuum of hybrid states according to their non-Pro-B RF classification probability (**Figure 2H**). This revealed transcriptionally hybrid populations with underlying ProB-like gene-expression, co-expressed with either progenitor-like genes (HSC-hyb) or genes implicated in later developmental phenotypes (PreB-hyb or ImmatureB-hyb) (**Figures S6A-C**). Genes correlated with these prediction probabilities reflected markers of earlier and later stages of B cell development (**Figures S5E & S6A; Table S5**), and largely agreed with our unbiased NMF results (**Figures S3A & S3C**). All three hybrid populations were characterized by predicted utilization of canonical transcription factors (TFs) active in the healthy reference cell subsets (e.g., CREB1, MYC in HSC-hyb; E2F2, FOXM1 in PreB-hyb; IRF4, FOXO3, CIITA in ImmatureB-hyb), as well as aberrant TF activity (e.g., IRF1, STAT1 in ImmatureB-hyb; **Figure S6D;** see **Methods**). Thus, anomalous co-expression of stage-associated genes in both primary patient samples and PDX models defines a hybrid development-like continuum in Ph+ ALL and implicates promiscuous, but still coherent, developmental transcriptional states.

### Hybrid development states are associated with treatment response and restricted mutation acquisition

We next asked whether shifts in this hybrid development-like continuum associated with resistance to combination TKI. Overall, progression samples were characterized by decreased hybrid population diversity, suggesting a restriction toward a single hybrid state (**Figure 3A**). Differential expression analysis across all PDX tumors revealed genes included in the ProB-like (e.g., *SOCS2, DNTT*) and HSC-hyb (e.g., *CD34, ID2, CD99*) signatures enriched at pre-treatment while genes implicated in the more mature PreB-hyb (e.g., *TCL1A, VPREB3, IGLL1*) and ImmatureB-hyb (e.g., *MS4A1, CD74, HLA-DRB1*) signatures were up-regulated at progression, implicating a shift into later developmental stages (**Figure 3B**). However, not all PDX models shifted toward more mature hybrid transcriptional states at progression. DFAB-25157, which progressed with mutations in *ABL1* (**Figure S2C; Table S3**), remained dominated by ProB-like and HSC-hyb states at both pretreatment and progression compared to other PDX lines (**Figures 3C & S7A**). Leukemias that progressed with RAS pathway mutations either contained a majority of cells expressing PreB-hyb and ImmatureB-hyb signatures at both pre-treatment and progression (CBAB-30198, DFAB-54880), or increased proportions of malignant cells with high PreB- and ImmatureB-hyb gene expression at progression (DFAB-62208; **Figure S2C; Table S3**). Notably, the two PDX lines that progressed with neither ABL nor RAS pathway mutations (CBAB-75914, CBAB-12402; **Figure S2C; Table S3**) demonstrated the strongest shifts toward more mature hybrid developmental bins.

This enrichment for more mature phenotypes at progression was a strong departure from patterns seen in Ph+ ALL treated with chemotherapy^11^ or imatinib,^8^ where progression on therapy was driven by less mature or stem-like cells. We sought corroborating evidence for this observation in our PDX trial samples using standard immunophenotyping approaches (**Figure S7B;** see **Methods**). Mirroring the transcriptional data, most pre-treatment leukemias harbored multiple subpopulations across the B cell developmental trajectory and showed a similar restriction in developmental state diversity at progression (**Figures 3D & S7C**). These immunophenotyping data also corroborated the overall enrichment of more developmentally mature phenotypes at progression (**Figure S7C**), specifically the predominance of more mature CD34-negative developmental phenotypes in leukemias that progressed with RAS pathway mutations or no mutations (*p*<0.001 from Dirichlet regression for both mutation group comparisons to ABL pathway-mutated leukemias; **Figures 3E, 3F & S7D**).

We next sought direct clinical evidence for the relevance of developmentally-hybrid programs in resistance to combination TKI. We prospectively collected serial single-cell measurements from the bone marrow of 2 patients (n=5 individual samples, n=7,649 cells; **Figure 3G; Table S6**) enrolled on a phase 1 trial testing dasatinib in combination with asciminib and prednisone. Clinical activity was assessed by the reduction in bone marrow *BCR::ABL1* mRNA transcript levels after three cycles of treatment (day 85; NCT02081378). Samples from patient BIAB-16768 maintained a predominant population of ProB-like malignant cells over the course of treatment and entered remission before 85 days of treatment (3-log reduction in bone marrow *BCR::ABL1* detected by qRT-PCR). By contrast, samples from patient DFAB-71417 rapidly shifted toward later developmental hybrid states (PreB-hyb and ImmatureB-hyb) by day 28 on therapy and failed to respond by day 85 (1-log reduction in bone marrow *BCR::ABL1* detected by qRT-PCR; **Figures 3G & 3H)**. Combined with our PDX analysis, these results provide preliminary evidence that more mature developmentally-hybrid expression programs can drive resistance to dual ABL1 inhibition.

**Figure 3.**
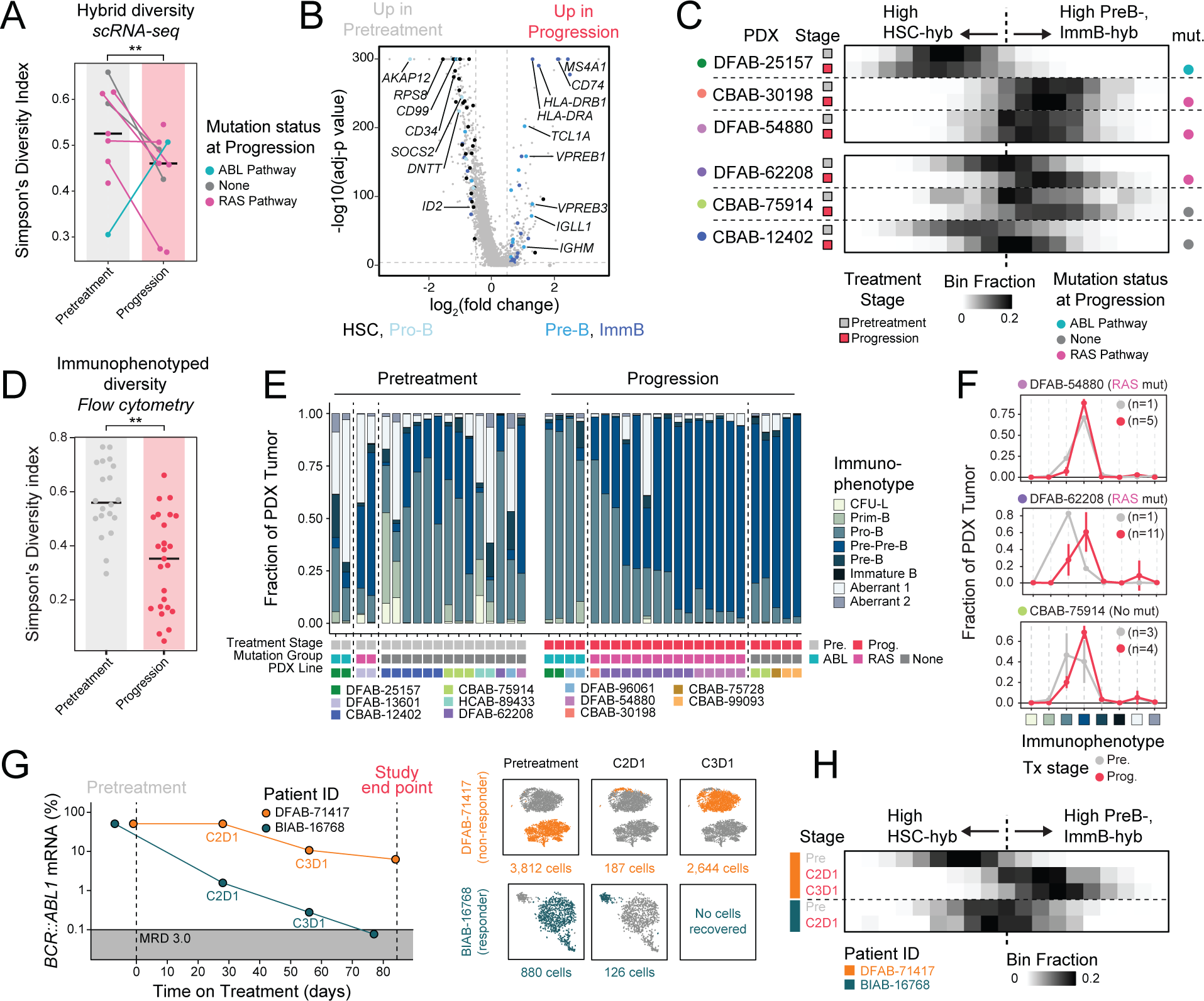
Oncogene withdrawal drives convergence onto developmental hybrids. **(A)** Simpson’s Diversity Index (SDI) of non ProB-like hybrid population proportions in each PDX line, colored by mutation status at progression. Tied points represent paired PDX treatment stages. Median SDI for pretreatment and progression across PDX lines plotted as a line. Wilcoxon rank sum p-value (\*\**p*<0.01) reported, excluding ABL pathway mutated PDX line (outlier DFAB-25157). **(B)** Differentially expressed genes between PDX pretreatment and progression single-cells. Marker genes for HSC, Pro-B, Pre-B, and Immature B cell types are annotated. **(C)** Density of cells across the spectrum of hybrid developmental gene expression space, calculated by the difference between later-stage hybrid scores (PreB-hyb, ImmatureB-hyb) and progenitor hybrid scores (HSC-hyb). Rows are annotated by PDX line, time point, and mutation (“mut.”) status at progression. **(D)** SDI of flow cytometry immunophenotyped B cell lineage populations within individual PDX tumors at pretreatment and progression; median SDI indicated for pretreatment and progression tumors. \*\**p*<0.01 (Wilcoxon rank sum test). **(E)** Fractional representation of immunophenotyped B cell lineage populations for 42 leukemia samples from 11 PDX lines at pretreatment and progression time points. “Pre.” = pretreatment; “Prog.” = progression. Immunophenotyped population flow cytometry markers defined in Figures S7B & S7C. **(F)** Pretreatment and progression average immunophenotyped population proportions (as plotted in (E)) for three representative PDX lines corroborate transcriptional trends in (C); error bars indicate ±1 standard deviation when at least 3 mice were profiled. Number of mice profiled at each time point indicated for each PDX line. PDX lines are labeled based on mutation group at progression. **G)** *BCR::ABL1* percent mRNA qRT-PCR traces (log10(BCR::ABL/β-Actin mRNA)) from bone marrow aspirates of two patients on combination ABL1 inhibition, including one representative responder (BIAB-16768) and one non-responder (DFAB-71417), from a Phase I clinical trial (**Table S6**). MRD 3.0 indicates trial definition of remission tumor burden (3-log reduction in bone marrow *BCR::ABL1* mRNA detected by qRT-PCR). Right: scRNA-seq data collected from patients at each treatment cycle time point shown on t-SNE projections. **(H)** Density of cells across the spectrum of hybrid developmental space, as defined in **(C)**, compared across paired patient pre-treatment and on-treatment time point bone marrow aspirates. *See also Figure S7; Table S6*.

### Longitudinal monitoring of cell state and mutational co-evolution

Collectively, our data nominate 3 potential routes of resistance to ABL1 inhibition in Ph+ ALL: 1) mutational reactivation of ABL signaling in progenitor-like states, 2) mutational activation of RAS signaling in later-stage hybrid states, or 3) transcriptional shifts toward later developmental hybrid states without accompanying mutational alterations. To directly explore whether these routes are recoverable at multiple timepoints during ABL1 inhibition, we next examined genotype-phenotype co-evolution by profiling single cells from pre-treatment, MRD (21 days on therapy), and progression in our PDX models, selecting individual leukemias that represent each putative mechanism of resistance (**Figure 4A**; DFAB-25157, ABL1 reactivation; DFAB-62208, RAS activation; CBAB-12402, no mutations). At each stage of therapy, we profiled leukemia cells using SMART-Seq2 (SS2)-based scRNA-seq to increase information capture from low cell numbers at remission and to facilitate matched single nucleotide variant (SNV) detection in the same single cells. For these longitudinal studies, we treated mice with single agent ponatinib (see **Methods**) since it performed equivalently to combination TKI therapy (**Figure 1C**) and is directly relevant to treatment being used in patients.

**Figure 4.**
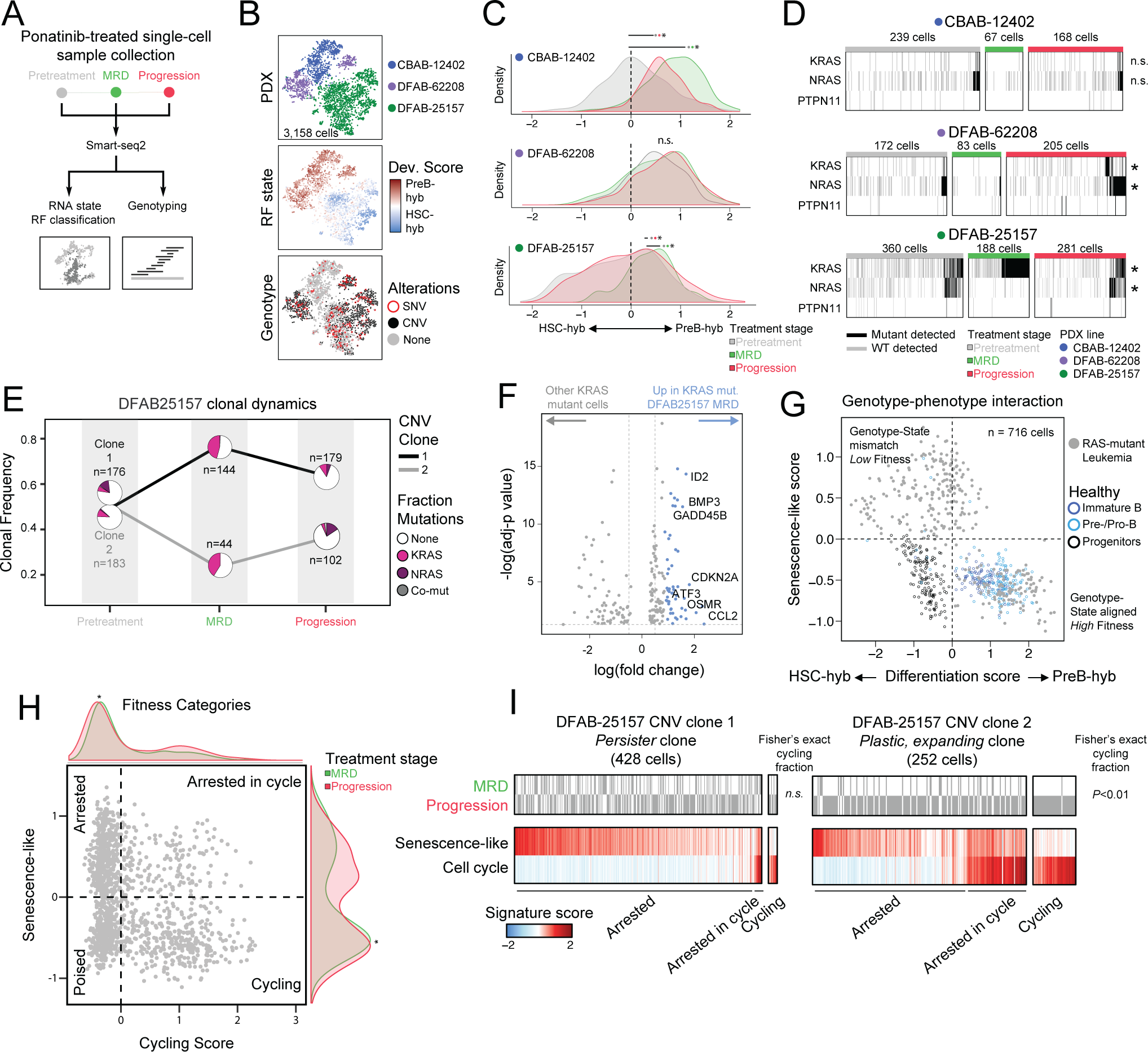
Developmental phenotypes restrict genotype fitness in remission. **(A)** Strategy for profiling three representative PDX models at pretreatment, MRD, and progression with Smart-Seq2 (SS2). **(B)** t-SNE visualizations for the leukemic cells collected with SS2 and labeled by PDX line (top), developmental state (middle; “dev.”=development), and detected genetic alterations (bottom; “SNV”=single nucleotide variant; “CNV”=copy number variant). **(C)** Density distributions of leukemia cells at pretreatment, MRD, and progression time points across HSC-hyb to PreB-hyb gene expression scores. *\*p*<0.001 from KS test for each pairwise comparison between treatment stages. **(D)** Mutant or wild-type (WT) transcript detection for *KRAS*, *NRAS*, and *PTPN11* within single-cells. Significant mutant transcript abundance between time points are annotated; \**p*<0.05 by Fisher exact test. **(E)** Dynamics of CNV sub-clonal proportions at pretreatment, MRD, and progression in DFAB-25157. Pie charts represent *KRAS* or *NRAS* fraction of each sub-clone at the indicated time points. Number of cells sampled within each CNV sub-clone are reported. **(F)** Differentially expressed genes between DFAB-25157 *KRAS*-mutant cells at MRD versus all other *KRAS*-mutant cells, highlighting increased expression of genes implicated in senescence (Table S8). **(G)** RAS-pathway mutant leukemic cells plotted according to their differentiation gene expression score on the x-axis, and senescence-like gene expression score on the y-axis. Overlaid healthy progenitor, Pre-B, and Immature B cells colored by cell type. **(H)** Fitness landscape of cell-cycle arrested, poised, and actively cycling leukemic cells in remission. Single-cells plotted by cycling (x-axis) and senescence (y-axis) signature scores. Distributions for cells in each fitness quadrant shown (green=MRD, red=Progression; \**p*<0.001 reported from KS test). **(I)** DFAB-25157 leukemic cells from each CNV subclone ranked along senescence-like and cell cycle signature scores. Fisher’s exact test p-value reported for the origin of cycling cells (MRD vs. Progression). No cells belonged to the “poised” fitness category from either CNV subclone. *See also Figures S8 & S9; Tables S7 & S8*.

First, we ensured the robustness of our RF hematopoietic developmental classifier on full-length, SS2 transcriptomes from both healthy (n = 421; same donors as **Figure S4**) and leukemic cells (n = 3,641; **Figure S8A;** see **Methods**). Using our RF framework, we independently derived the leukemic cellular states in our SS2 dataset (**Figures S8B-D; Table S7**), finding they highly correlated with our Seq-Well-derived hybrid phenotypes – specifically in early progenitor (Progenitor-like vs. HSC-hyb) and more mature (PreB-like vs. PreB-hyb and ImmatureB-hyb) leukemic cell states (**Figure S8E**). Given this coherence, hereafter we refer to Progenitor-like and PreB-like SS2 programs as HSC-hyb and PreB-hyb respectively for simplicity. We next detected mutated transcripts identified from bulk DNA sequencing within individual cells from our SS2 data (**Figure S9A; Table S3;** see **Methods**). The number of detected mutant transcripts in SS2 libraries was limited by the average expression of the corresponding gene, with higher rates of detection for RAS pathway single-nucleotide variants (SNVs; *GNB1, NRAS, KRAS, PTPN11*) compared to ABL pathway SNVs (*ABL1, STAT5A*) (**Figure S9B**). For highly expressed target genes, however, the proportion of single cells harboring mutations corresponded with the variant allele frequency measured in bulk sequencing of the same tumor (**Figure S9C; Table S3**), highlighting that SS2 provides sufficient SNV detection to capture the kinetics of RAS pathway mutations in our dataset. Furthermore, single-cell profiling enabled highly sensitive detection of rare malignant cells harboring mutations with less than 3% VAF from bulk sequencing, allowing comparisons of dominant and rare subclones (**Figure S9C**). Finally, we identified copy number variations (CNVs) in the SS2 profiles using inferCNV (see **Methods**). In combination with transcriptional state information, these data provided a detailed, high-resolution picture of the co-evolution of mutational and transcriptional heterogeneity in B-ALL single cells over the course of ponatinib treatment (**Figures 4B & S9D-F**).

### Cell state dictates fitness and restricts growth of RAS-mutant cells in remission

Using this high-resolution dataset, we first evaluated changes in hybrid developmental state frequency between pre-treatment and residual cells in each model during treatment with ponatinib (**Figure 4C**). CBAB-12402 was transcriptionally dynamic and demonstrated a significant shift towards a dominant PreB-hyb phenotype among MRD cells that was conserved at progression, mirroring patterns seen in the larger PDX trial for this model (**Figures 3C & S9F**). Transcriptional states in MRD and progression leukemia cells from DFAB-62208 displayed a minor shift forward to stronger PreB-hyb expression compared to pre-treatment. DFAB-25157 was variable along the progenitor to mature phenotype continuum at both pretreatment and progression, driven by dominant HSC-hyb gene expression. A subset of cells from this model co-expressed HSC-hyb and PreB-hyb states in MRD, albeit at much lower levels than PreB-hyb scores in the other two models (**Figures S9E-G**).

Point mutations in *NRAS* and *KRAS* from the same leukemia cells revealed surprising dynamics across PDX models and stages of therapy (**Figure 4D**). We detected very low frequency RAS mutations in CBAB-12402 at pre-treatment that were not enriched at progression, in agreement with bulk DNA sequencing data that did not identify actionable driver mutations (**Figure S2C; Table S3**), thus implicating a “state-shift” only mechanism enabling progression. DFAB-62208 also harbored low-frequency *KRAS* and *NRAS* point mutations at pretreatment; a single *NRAS*-mutant, cycling cell was observed in remission and both *KRAS*- and *NRAS*-mutant clones expanded at progression (mirroring bulk sequencing data; **Figure S2C; Table S3**), suggesting the preexisting PreB-hyb transcriptional state was permissive for expansion of RAS-mutant clones. In DFAB-25157, we observed a significant increase in the proportion of *KRAS* mutant malignant cells in MRD (3 of 6 mice at MRD harbored identifiable RAS-mutant cells) compared to pretreatment leukemic cells, a finding we confirmed using bulk DNA sequencing from a separate sample (**Table S3**; Mouse 4H0, *KRAS* AF 0.75). This was surprising given that this model does not progress on therapy with emergent RAS mutations (**Figures 4D & S2C**). Indeed, considering both single-cell CNV and SNV clones (**Figures S9D & S9E**), we found no evidence of outright genetically-driven clonal selection in DFAB-25157 despite the enrichment of RAS-mutant cells in remission (**Figure 4E**). In this case, our data suggest that RAS-family mutations in cells with a discordant developmental cell state permit survival (or persistence) in the context of ABL inhibition but confer a fitness disadvantage that suppresses their expansion.^10^

We next interrogated the single-cell transcriptomes of remission DFAB-25157 cells to define mechanisms for this apparent state-genotype incompatibility. *KRAS*-mutant leukemic cells from DFAB-25157 at MRD upregulated genes that positively regulate senescence (e.g., *CCL2, TOB1*) and negatively regulate cell cycle (e.g., *CDKN2A*) compared to *KRAS*-mutant leukemia cells from all other time points and PDX lines (**Figure 4F**). To evaluate how this signature evolves over the course of therapy, we scored individual cells for these upregulated senescence-associated genes (Senescence-like score; **Table S8**). *KRAS*-mutant clones with similar senescence-like signatures were present at pretreatment in cells with co-incident HSC-hyb phenotypes, whereas PreB-hyb *KRAS*-mutant leukemia cells across other treatment stages and PDX lines had low senescence-like scores (**Figure S9G**). These data suggest the fitness of RAS mutant clones is influenced by the compatibility of transcriptional state and genotype: the expression of senescence-implicated genes is restricted to HSC-hyb cells harboring RAS mutations, whereas RAS-mutant PreB-hyb cells remain capable of entering the cell cycle (**Figure 4G**). Therefore, despite activation of a mitogenic oncogene that contributes to resistance to TKI in multiple contexts, developmental states restrict the expansion of these genotypes, including during deep remissions.

As MRD genotypes alone could not predict clonal expansion driving progression, we sought to identify what phenotypes persist in MRD and actively contribute to progression. We binned each cell from MRD and progression into four fitness phenotypes based on their expression of senescence-like and cell cycle scores (**Figure 4H**). To our surprise, progression contained a significant accumulation of putatively cell cycle-arrested cells with higher senescence-like scores compared to MRD (*p*<0.001, KS statistic). Notably, we also observed CNV subclonal fitness plasticity in DFAB-25157, whose cells at MRD were characterized by high senescence-like scores. A cycling population of RAS-wildtype cells from one subclone emerged at progression (**Figures 4I & S9H;** *p*<0.01, Fisher’s exact test), associated with an increased abundance of that subclone at progression (**Figure 4E**). In contrast, RAS-mutant cells from DFAB-62208, characterized by later developmental phenotypes, were highly proliferative at progression (**Figure S9H**). Collectively, these data suggest that diverse Ph+ ALL genetic subclones can persist to progression and even clones with senescence-like phenotypes at MRD may expand with enhanced fitness to seed progression. Given the possibility of plasticity and the restrictions imposed by cell states on certain genotypes, these data suggest it may be difficult to predict from genetics alone the subclones that will ultimately seed relapse.

### Direct targeting of transcriptional programs in residual disease deepens remission

In light of this complexity, we hypothesized that directly targeting transcriptional programs that enable persistence at MRD could overcome the diversity of subclones identified at remission. Using differential expression and gene-gene correlation (see **Methods**), we identified three expression programs in remission that persisted to progression – a Pre-BCR Signaling program, closely aligned with the PreB-hyb state (e.g., *IGLL1, VPREB3*), a Stress/Autophagy program (e.g., *HSPA1A, UBC*), and an inflammatory program (e.g., *EGR1, JUN, TNF*; **Figure 5A; Table S9**). The inflammatory program was evenly expressed across all leukemic cells in remission, a phenotype seen in other hematological diseases (28504724, 35618837; **Figure S10A**). The remaining expression programs were variable across MRD cells stratifying those high for the Stress/Autophagy cell state and those expressing the Pre-BCR Signaling program (**Figures S10A & S10B)**. We considered these variable programs to test the hypothesis that targeting specific expression programs could deepen remissions. These two variable gene expression programs split along fitness subpopulations, with leukemic cells harboring high Pre-BCR Signaling scores also scoring high for cell cycle, and leukemic cells with high Stress/Autophagy program scores enriched for senescence-like expression (**Figure 5B**).

**Figure 5.**
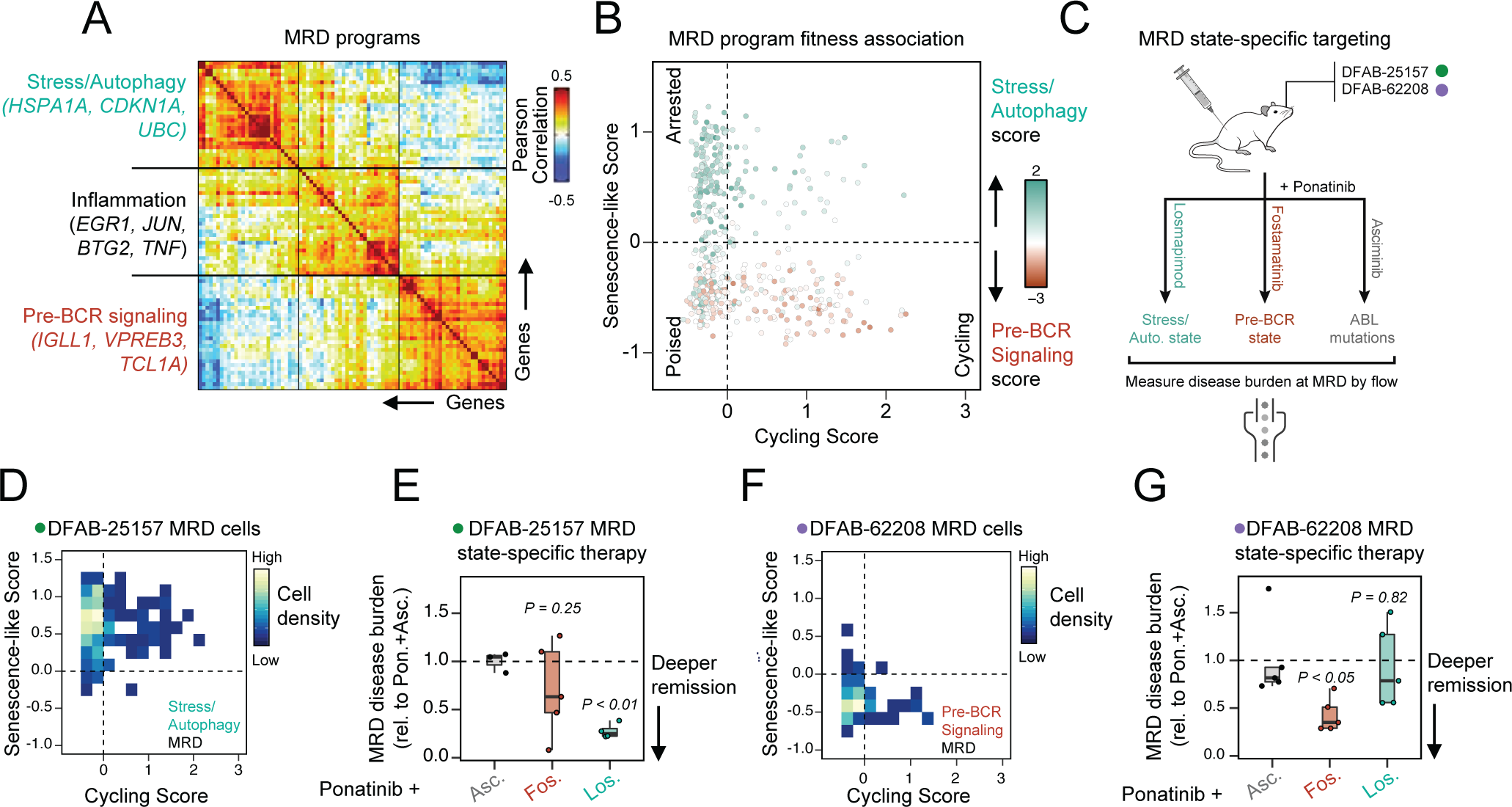
Targeting integrative cell states enhances remission. **(A)** Pairwise Pearson correlation of genes defining MRD states (Table S9). **(B)** Module scores for the Stress-Autophagy (turquoise) and Pre-BCR Signaling (dark red) states projected over single-cells at MRD. Cells are plotted along fitness quadrants as in **Figure 4H** by their cycling (x-axis) and senescence-like (y-axis) gene signature scores. **(C)** Study design for testing MRD cell-state targeting. **(D)** Cell fitness distribution for DFAB-25157 MRD cells. **(E)** DFAB-25157 MRD bone marrow disease burden assessed by flow cytometry (y-axis, relative to Ponatinib+Asciminib) in the respective treatment arms (“Asc.”=Asciminib; “Fos.”=Fostamatinib; “Los.”=Losmapimod). T-test p-values reported, comparing losmapimod and fostamatinib arms to asciminib reference. **(F)** Cell fitness distribution for DFAB-62208 MRD cells. **(G)** DFAB-62208 MRD disease burden assessed by flow cytometry as in **(E)**. Reported t-test p-values compare losmapimod and fostamatinib arms to asciminib reference. *See also Figure S10; Table S9*.

We next evaluated whether these two gene expression programs could be therapeutically targeted. We paired ponatinib with either the FDA-approved SYK inhibitor, fostamatinib, to inhibit pre-BCR signaling in leukemic cells scoring highly for the Pre-BCR program, or the FDA-approved p38a MAPK inhibitor losmapimod, to target leukemic cells scoring highly for the Stress/Autophagy program given the co-enrichment of p38a MAPK activation with the Stress/Autophagy program and previous work supporting crosstalk between p38 signaling and autophagy/leukemic stem cell-related phenotypes (**Figure S10C**).^33,34,35^ As a combination control, we compared transcriptional-state-directed combination therapy to dual oncogene targeting using ponatinib and asciminib (**Figure 5C**). We selected two PDX lines that were enriched for either variable MRD expression program: DFAB-25157, which scored highly for the Stress/Autophagy program, and DFAB-62208, which scored highly for the Pre-BCR signaling program and sat along the poised/cell cycle spectrum (**Figures 5D, 5F & S10D**). DFAB-25157 mice treated with combination losmapimod plus ponatinib showed a significant reduction in residual disease burden compared to dual oncogene suppression, a striking comparison as DFAB-25157 tumors consistently progressed with acquired mutations in ABL1 (**Figures 5E & S2C**). Analogously, DFAB-62208 mice responded to ponatinib plus fostamatinib and had significantly reduced residual disease compared to dual oncogene suppression (**Figure 5G**). These data suggest that residual leukemia cells can be effectively targeted according to the specific transcriptional state governing persistence in remission.

### A biophysical workflow for low-cost, rapid coupling of genotype to developmental state in leukemia cells

Our data support the importance of both mutations and overall cell state in determining leukemic cell fitness and therapeutic susceptibility at MRD. While mutations can be monitored in clinical workflows from residual leukemic cells, single-cell transcriptomics is currently difficult to scale due to the overall cost and time required for sample collection and analysis. We sought a metric that would integrate complex transcriptional information from low-input MRD samples to enable rapid determination of leukemic cell state, compatible with downstream mutational profiling.

Immunophenotyping strategies of developmental cell states, especially given the very low cell numbers at MRD, is likely to be highly challenging. Alternatively, cell size characteristically decreases as healthy progenitor cells progress from HSCs to pro-B to pre-B cells, putatively providing a label-free attribute with which to phenotype ALL cells.^36^ We have previously shown that measurements of buoyant mass, as measured by the suspended microchannel resonator (SMR),^37^ can reveal changes in cell state.^38,39,40,41,42^ Buoyant mass (referred to hereafter simply as mass) can be measured from live single cells with a resolution near 50 fg, which is highly precise given that the average buoyant mass of a hematopoietic cell is ∼75 pg.^43^ Further, we have shown that coupling mass measurements to scRNA-seq from the same cell enables the determination of expression-dependent changes in cellular mass.^41^ Thus, we hypothesized that underlying biophysical development-like phenotypes may be conserved and sufficient to rapidly capture the developmental state of a leukemia cell.

We first determined whether mass can distinguish B cell developmental states in healthy donors. By performing paired SMR-SS2^41^ on cells flow-sorted from healthy donors into Progenitor (CFU-L; 155 cells), Pro-B (122 cells), and Immature B (105 cells) gates, we found that each stage of B cell development was characterized by distinct mass distributions, with decreasing cell mass along the B cell developmental trajectory (**Figures 6A, 6B, S11A & S11B**). Within each B cell developmental stage, healthy cells with higher mass also scored highly for S phase or G2/M phase cell cycle, a pattern seen across studies using SMRs within a specific cell type (**Figure S11A**).^41,43,44^ We found a strong relationship between each gene’s dependence on RF prediction scores and matched cellular mass (r = 0.88 from Pearson correlation), indicating that genes highly associated with cell mass are also most correlated to healthy B cell developmental states (**Figure S11C**). Consistently, in leukemic cells, genes defining the HSC-hyb signature were most positively correlated with leukemic cell mass, and genes defining the PreB-hyb signature were most negatively correlated with cell mass (r = 0.90) (**Figure 6C**). We validated this observation across 17 additional PDX samples at the bulk level showing that the average leukemic cell mass reflects the average RF predicted state (r = 0.66) and tracks with the progression-emergent mutations for each PDX (**Figure 6D**). Taken together, these data support mass as a meaningful surrogate for development-associated transcriptional state in leukemia cells.

**Figure 6.**
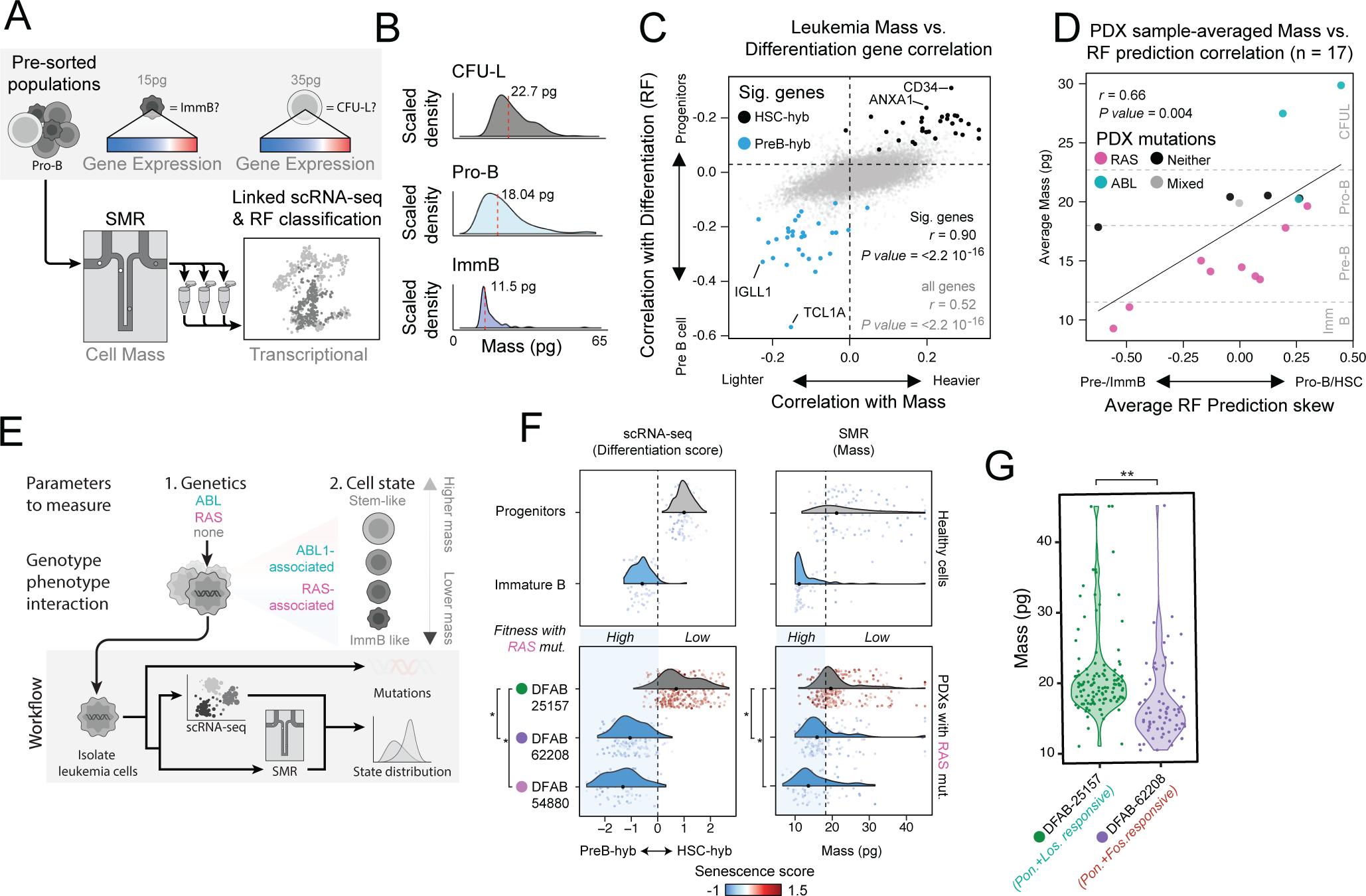
Biophysical measurements can be used as a surrogate for complex transcriptional states. **(A)** Schematic for evaluating the relationship between complex transcriptional state and integrative biophysical features. **(B)** Mass distributions from the sorted populations in (A) measured using the SMR; median mass reported. **(C)** Leukemia cell mass-correlated genes (x-axis) are plotted against each gene’s correlation to developmental phenotypes (RF probability for progenitor and Pre-B cell types; y-axis). Colored points mark genes included in the Progenitor and Pre-B SS2 signatures; “Sig. genes”=Leukemia developmental marker genes. **(D)** Average difference in RF prediction score between early and late stages of B cell development (x-axis) versus average mass for each mouse (n=17), binned by distributions in (B) and Figure S11A, and annotated by progression mutation status. **(E)** Proposed workflow for comparing sequencing to biophysical measurements for diagnostics. **(F)** Example application for pairing mutation and mass information to predict development and fitness-integrated transcriptomic state. Density spectra of (left) developmental score and (right) mass for (top) healthy progenitor cells and immature B cells, and (bottom) RAS-mutant leukemic cells in three representative PDX lines. Dotted line for mass distribution represents mean+1 standard deviation of healthy Immature B mass. Median differentiation scores or mass for each PDX line are denoted as a dot; PDX lines are colored based on their median similarity to Immature B or Progenitor differentiation scores or mass. * indicates significant difference between DFAB-25157 differentiation score or mass distributions compared to those of DFAB-62208 and DFAB-54880 (KS test, *p*<0.001). Individual cells are colored according to their senescence signature score. Blue shaded region is the putative zone of compatibility for RAS mutations and developmental state. **(G)** Mass distributions for leukemia cells at MRD from DFAB-25157 (sensitive to combination losmapimod) and DFAB-62208 (sensitive to combination fostamatinib). \*\**p*<0.001 from paired Wilcoxon test. *See also Figure S11*.

Finally, we evaluated how single-cell mass could pair with genotyping to further define developmental state and mutation compatibility (**Figure 6E**). We compared mass distributions between RAS-mutant PDX lines with higher HSC-hyb and high senescence-like gene expression (DFAB-25157) and PDX lines with higher PreB-hyb gene expression (DFAB-62208 and DFAB-54880). State-genotype discordant HSC-hyb DFAB-25157 cells were enriched for senescent-like scores and significantly higher mass than the more developmentally-mature and non-senescent DFAB-62208 and DFAB-54880, mirroring mass differences between healthy progenitor and immature B cells (**Figure 6F**). Furthermore, we found a significant difference between the mass distributions of DFAB-25157 MRD cells compared to DFAB-62208 cells at MRD (**Figure 6G**), implicating that mass measurements reflect developmentally-relevant and therapeutically actionable heterogeneity in MRD for these leukemias (**Figures 5D-G**). Consequently, mass measurements appear to be sufficiently sensitive to distinguish differences in developmental state for leukemic cells, and, when assessed simultaneously with genotypic data from the same sample, may predict therapeutic susceptibility for targeting states in MRD.

## DISCUSSION

Oncogene-directed therapy provides clear benefits to certain patient populations, yet it is equally clear that targeting cancers solely based on their mutational heterogeneity has an upper limit.^45,46,47^ Indeed, our phase II-like preclinical trial results reveal that even combinations of highly potent TKIs aimed at the same oncogene do not cure Ph+ ALL. While much of the preclinical and clinical data in CML and ALL have identified pathway reactivation through alterations in ABL1 as a primary mechanism of escape,^1,2,3,4^ our data suggest alternative pathway activation through RAS alterations also drives resistance in a significant fraction of cases. Mirroring patterns seen in patients,^5^ our trial also shows that a large fraction of mice engrafted with patient-derived leukemias (up to 40%) progress without a clear genetic driver, warranting the exploration of alternate therapeutic strategies for these cases.

Transcriptional phenotypes have been described in AML,^16^ CML,^13^ and ALL,^8^ and recent studies suggest that patients with more progenitor-like leukemia cells have a worse overall prognosis and tend to respond poorly to therapy. In ALL specifically, a recent study showed that leukemias enriched for progenitor-like states have worse outcomes on imatinib.^8^ Our data suggest that lineage plasticity is relatively common in response to 3^rd^ generation and combination TKI therapy, with resistant leukemia cells most frequently mimicking later stages of B cell development. This contrasts with most settings where, even in solid cancers, a canonical response to therapy is the enrichment of less differentiated cell states.^48,49^ Moreover, we demonstrate the importance of defining cell state and mutational associations – despite myriad mutational routes that might be predicted to confer resistance, our data suggest that specific transcriptional backgrounds may restrict leukemias to distinct subsets of escape mutations. Though these associations will need to be learned in larger cohorts and for each specific disease, this framework may represent a strategy for prioritizing the permissible transcriptional state/mutational convergences within oligo/polyclonal populations that can drive progression.

While there is agreement on the clinical and therapeutic importance of understanding MRD, the phenotypes of the residual cells responsible for seeding progression and how to best target them remains an outstanding question in the field owing to several technical challenges.^21,25^ In this regard, Ph+ ALL is a tractable system, as it is feasible to isolate MRD from either blood or bone marrow of patients or xenografted mice in adequate numbers to allow for single-cell transcriptomics in addition to DNA sequencing. We found that matched genotypic and phenotypic profiling of rare MRD cells was critical for identifying three key insights about the biology of MRD and the translational potential of targeting it prior to relapse. First, the conventional wisdom proposes that not all cells at MRD can seed relapse, especially those that have exited the cell cycle or are otherwise classified as “unfit”.^21,25^ In contrast, we find that some CNV-defined clones expressing senescence-like genes at MRD can re-enter the cell cycle and contribute to progression. Of note, a similar phenotype has also been observed in AML treated with chemotherapy.^50^ Second, our discovery that senescent clones harboring RAS mutations were enriched in residual disease but did not contribute to relapse highlights the importance of understanding the cell state of mutant cells. This observation complicates current MRD evaluation strategies, as information about genotype alone will likely be insufficient to predict relapse for specific leukemias. Third, we show that co-targeting tumor-specific transcriptional programs in remission out-performs additional targeting of the same oncogene, at least with current therapeutics. This finding provides a translational rationale for identifying transcriptional phenotypes in residual disease to inform the rational selection of combination strategies. The importance of targeting cell state likely extends to other cancers where a central oncogene can be deeply inhibited, resulting in relapses that have acquired an alternate histology, including small cell relapse after androgen receptor inhibition in prostate cancer,^51^ squamous cell and small cell transitions after EGFR inhibition in lung adenocarcinoma,^52,53^ and estrogen receptor positive relapse after HER2 blockade.^54^

We note that the influence of an intact immune system on the developmental dynamics of Ph+ ALL is not well defined and represents a liability of our approach interrogating PDX models of leukemia in NSG hosts. We mitigated this by confirming our PDX results in serial measurements from patient bone marrow, but future efforts should include the use of humanized xenograft models and additional evaluation of primary patient specimens. Nevertheless, our identification of a central role for developmental state in Ph+ ALL has had immediate clinical implications. Our phase 1 clinical trial of dual oncogene targeting (NCT03595917) completed accrual^55^ and reopened as a phase 2 trial incorporating early introduction of the CD3xCD10 bispecific antibody blinatumomab (anti-CD3xCD19 bispecific antibody), which should maintain activity across the developmental states we have defined in MRD and relapse. Importantly, blinatumomab has demonstrated promising clinical activity in clearing residual disease in patients intended for consolidative allogeneic hematopoietic stem cell transplantation.^56,57^

Evaluating complex, non-mutational biomarkers may have significant clinical challenges. scRNA-seq is not yet a clinically-scalable assay, nor is it readily interpretable on a short time-scale. For translation to clinical workflows, it will be critical to develop diagnostics that are able to assess a sample’s genotype and relevant phenotype with reasonable throughput and interpretability. For remission profiling specifically, this is further complicated by the requirement for use with low-input samples. Owing to the low-input and non-destructive nature of the SS2-SMR measurement,^41^ we were able to acquire a unique dataset that directly links cellular mass to leukemic developmental state. These data establish that assessing complex, non-mutational biomarkers may be possible using mass as a relatively simple integrative cellular property. Our matched SMR/scRNA-seq data from normal bone marrow hints that mass variation may extend to other hematopoietic lineages as well so this approach may be applicable in diseases with significant developmental heterogeneity such as AML.^16^ We speculate that additional features of clinical utility in different disease contexts could come from other integrative single-cell properties such as morphology.^58^

In sum, we find transcriptional state controls the fitness of individual clones in MRD and dictates the landscape of progression on TKI in Ph+ ALL. We highlight the need to understand and monitor both mutational and transcriptional features in clinical pipelines to properly evaluate individual clones for their potential to drive relapse. We functionally establish the paramount importance of cell state in this context and suggest it should be prioritized for targeting in conjunction with driver oncogenes. In agreement with recent studies in solid cancers,^59,60,61^ our work in leukemia makes it apparent that therapies intended to convert remissions to cures should consider monitoring and targeting features outside of traditional mutational biomarkers.^62^

## ACKNOWLEDGEMENTS

This work was funded by the NIH-NCI U54 CA217377 (S.R.M., D.M.W., A.K.S.), K08 CA212252 (M.A.M.), K12 HL141953-05 (M.A.M.), P30 CA14051 (A.K.S., S.R.M.), 1U2C CA23319501 (A.K.S.), R35 CA231958 (D.M.W.); the Paul G. Allen Frontiers Group Distinguished Investigator Award (S.R.M., D.M.W.); the Sloan Research Fellowship in Chemistry (A.K.S.); and the Pew-Stewart Scholars Program for Cancer Research (A.K.S.). The authors acknowledge assistance with targeted panel and whole exome sequencing of PDX specimens from Dr. Aaron Thorner, Dr. Anwesha Nag, and Neil Patel of the Dana-Farber Cancer Institute Center for Cancer Genomics.

## AUTHOR CONTRIBUTIONS

Conceptualization, P.S.W., S.R.M., D.M.W., A.K.S. and M.A.M.; Methodology, P.S.W., M.L.R., A.W.N., S.S., C.P.C., L.C., S.R.M., D.M.W., A.K.S., and M.A.M; Validation, P.S.W., M.L.R., A.W.N., and M.A.M.; Formal Analysis, P.S.W., M.L.R., K.E.S., S.R., A.D., S.S., and M.A.M.; Investigation, P.S.W., M.L.R., A.W.N., A.D., S.S, H.S., N.S., M.M., H.H.A., L.B., P.D., C.S.L., K.S., J.G.R., Y.Z., F.P., N.M., L.C., A.P.A, S.V.R., A.J.G., N.C., A.V.S., K.J., H.L., R.J.K., M.M.S., M.A.M.; Resources, S.R.M., D.M.W., A.K.S., and M.A.M.; Data Curation, P.S.W., A.W.N and M.A.M.; Writing – Original Draft, P.S.W., M.L.R., A.W.N, A.K.S., and M.A.M; Writing – Review & Editing, P.S.W, M.L.R., A.W.N, L.C., A.P.A., S.S., S.V.R., M.R.L., S.R.M., D.M.W., A.K.S., and M.A.M.; Visualization, P.S.W., M.L.R., A.W.N., A.K.S., and M.A.M.; Supervision, M.A.M., P.S.W., S.R.M., D.M.W., and A.K.S.; Project Administration, P.S.W., S.R.M., D.M.W., A.K.S., and M.A.M.; Funding Acquisition, S.R.M., D.M.W., A.K.S., and M.A.M.

## METHODS

### RESOURCE AVAILABILITY

#### Lead Contact

Further information and requests for resources and reagents should be sent to and will be fulfilled by Dr. Peter Winter.

#### Data Availability

The scRNA-seq data and SMR data reported in this paper will be deposited in a central data sharing repository (Genomic Data Commons) under the NCBI Database of Genotypes and Phenotypes (dbGaP). scRNA-seq digital gene expression matrices, metadata, and interactive visualization tools will additionally be available through the Alexandria Project, a Bill & Melinda Gates Foundation-funded portal (part of the Single Cell Portal hosted by the Broad Institute of MIT and Harvard). Code used for analysis will be available upon request.

### EXPERIMENTAL MODEL AND SUBJECT DETAILS

#### Generation and Use of PDXs

Primary bone marrow and peripheral blood specimens were collected from patients with leukemia at the Dana-Farber Cancer Institute, Brigham and Women’s Hospital, and Boston Children’s Hospital for xenotransplantation. Additional PDXs that had already been established through the Public Repository of Xenografts (PRoXe) were utilized.^27^ De-identified patient samples were obtained with informed consent and xenografted under Dana-Farber/Harvard Cancer Center Institutional Review Board (IRB)-approved protocols. Nod.Cg-*Prkdc^scid^IL2rg^tm1Wjl^*/SzJ (NSG) mice were purchased from Jackson Laboratories and handled according to Dana-Farber Cancer Institute Institutional Animal Care and Use Committee-approved protocols. Salient PDX line metadata are provided in **Tables S1 & S2**.

#### *In vivo* therapeutic studies

Viably frozen Ph+ ALL xenograft cells were thawed and changed into 1X PBS before tail-vein injection at 0.5-2.0*10^6^ cells per mouse. Engraftment was monitored by weekly peripheral blood flow cytometry beginning three weeks after injection. Blood was processed with Red Blood Cell Lysis Buffer (Qiagen #158904; Hilden, Germany) and stained with antibodies against human CD45 (APC-conjugated, eBioscience #17-0459-42; San Diego, CA, USA) and human CD19 (PE-conjugated, eBioscience #12-0193-82) in 1X PBS with EDTA (2mM). Flow cytometry data were analyzed using FlowJo software (BD Biosciences; Ashland, OR, USA). Upon engraftment – when at least 10% of cells were positive for CD45 and CD19 – mice within each PDX line underwent 1:2:2:4:1 randomization to the following arms and initiated treatment within two days: (1) sacrifice for baseline tissue interrogation; (2) ponatinib (Selleckchem #S1490; Houston, TX, USA; constituted in 25mM citrate buffer, pH 2.75) 40mg/kg via oral gavage (OG) daily; (3) asciminib (NVP-ABL001, Novartis Pharmaceuticals; Basel, Switzerland; constituted in HCl 0.1M, PEG300 30%, Solutrol HS15 6%, NaOH 0.1M, sodium acetate buffer pH 4.7 10mM) 30mg/kg OG twice daily; (4) ponatinib 40mg/kg OG twice daily plus asciminib 30mg/kg OG BID; and (5) vehicle (alternating doses of vehicle used for ponatinib and asciminib, at equivalent volumes). One mouse per active treatment arm per PDX line was sacrificed on day 7 of treatment for pharmacodynamic assessment. The remaining mice continued daily treatment under monitoring with biweekly peripheral blood flow cytometry until progression (defined as peripheral blood involvement of at least 10% on two consecutive assessments at least one week apart), weight loss of greater than 20% from pre-treatment baseline, or clinical manifestations of advanced disease, including but not limited to ruffled fur, hunched posture, hind limb paralysis, or lethargy. Progression or toxicity as defined above triggered humane euthanasia by CO_2_ asphyxiation, necropsy to ascertain cause of death, and post-mortem harvest of peripheral blood, bone marrow, and any soft tissue masses. Additional *in vivo* studies involved treatment with nilotinib (Selleckchem #S1033), which was constituted in N-methyl-2-pyrrolidone (10%) in polyethylene glycol (PEG)-300 (90%) and dosed at 50mg/kg OG twice daily.

Studies to define the *in vivo* activity of combination therapies targeting the biology of MRD within individual PDX lines DFAB-62208 and DFAB-25157 utilized the same xenotransplantation and engraftment monitoring scheme as previously described and the following drugs: ponatinib (as above), asciminib (as above), fostamatinib (Selleckchem #S2206-50mg), constituted in 0.1% carboxymethylcellulose sodium, 0.1% methylparaben, and 0.02% propylparaben (pH 6.5) and dosed at 25mg/kg OG thrice daily, and losmapimod (Selleckchem #S7215-50mg), constituted in 1% DMSO in methylcellulose and dosed at 20mg/kg via the intraperitoneal (IP) route daily. Upon engraftment (>10% leukemia involvement of peripheral blood), individual mice underwent live femoral bone marrow aspirates under anesthesia with inhaled isoflurane delivered via precision vaporizer and underwent 1:1:1 randomization to the combination of ponatinib and asciminib, ponatinib and fostamatinib, or ponatinib and losmapimod. Animals initiated treatment within 48 hours of engraftment and continued treatment for 21 days ± 3 days, at which point they underwent humane euthanasia, necropsy, and immediate post-mortem recovery of peripheral blood and bone marrow from the femur contralateral to that which was aspirated upon engraftment.

#### Human donors for reference

Normal human bone marrow aspirates were obtained from donors who provided informed consent for tissue banking and research under Dana-Farber/Harvard Cancer Center IRB protocols and were undergoing bone marrow harvest for unrelated hematopoietic stem cell transplantation recipients. Briefly, bone marrow was collected into a Baxter bone marrow harvest collection system with diluent consisting of sodium heparin in lactated Ringers solution. Bone marrow was heparinized at a final concentration of 15-20 units/mL and filtered inline using 200μm and 500μm filters. Bone marrow mononuclear cells from the heparinized, filtered product were isolated via density gradient centrifugation (Ficoll-Paque, ThermoFisher Scientific #45-001-749) and subsequently underwent fluorescence-activated cell sorting (FACS) to isolate hematopoietic developmental subpopulations for Seq-Well S^3^ and SS2 single-cell transcriptomic profiling (see Methods Details).

#### Phase I clinical trial

Serial primary blood and bone marrow specimens were obtained from appropriately consented patients treated on a phase I, investigator-initiated clinical trial (NCT03595917) of asciminib (ABL001) in combination with dasatinib plus prednisone for adults with newly diagnosed Ph+ ALL or chronic myelogenous leukemia in lymphoid blast phase (CML-LBP). Some patients cross-consented to a Dana-Farber Cancer Institute tissue banking protocol permitting additional evaluation of primary specimens. Bone marrow was obtained at screening and after each 21-day cycle through the first four cycles. Peripheral blood was obtained at screening and on days 2, 4, 8, 11, 15, and 22 (±2 days) of cycle 1. Both bone marrow and peripheral blood were collected into EDTA vacutainer tubes prior to mononuclear cell isolation per standard protocols. Bone marrow and peripheral blood underwent clinical quantitative real time PCR for *BCR::ABL1* mRNA according to the *BCR::ABL1* isoform detected at screening (p190 or p210). Curated sets of Ph+ ALL clinically annotated specimens underwent evaluation by scRNA-seq (Seq-Well S^3^; salient donor metadata provided in **Table S6**).

## METHOD DETAILS

### Quantifying *BCR::ABL1* mRNA in PDX peripheral blood with qRT-PCR

*BCR::ABL1* mRNA levels were measured via quantitative real-time PCR (qRT-PCR) of serial peripheral blood specimens from PDX models to track kinetics of response and progression. Briefly, xenografted mice were phlebotomized for 100µL by submandibular vein laceration every two weeks. Blood was stored in RNAProtect tubes (Qiagen #76544). mRNA was isolated using the RNeasy Protect Animal Blood Kit (Qiagen #73224) and quantified using the iScript One-Step RT-PCR Kit with SYBR Green (Bio-Rad #170-8893) on a Bio-Rad CFX96 Thermal Cycler. Synthesis of cDNAs was performed with random hexamers. Amplification of cDNAs was performed using iTaq Universal SYBR Green Supermix (Bio-Rad #172-5125) and the following oligomers:

*BCR::ABL1* isoform p190 forward: CAACAGTCCTTCGACAGCAG

*BCR::ABL1* isoform p190 reverse: CCCTGAGGCTCAAAGTCAGA

*BCR::ABL1* isoform p210 forward: TCCGCTGACCATCAATAAGGA

*BCR::ABL1* isoform p210 reverse: CACTCAGACCCTGAGGCTCAA

Positive control reagents for each isoform were p190 clonal control RNA (Invivoscribe #4-089-2800) and mRNA isolated from the BCR::ABL1 p210-positive cell line K562.

### Quantifying *BCR::ABL1* mRNA in primary patient peripheral blood with qRT-PCR

*BCR::ABL1* mRNA was quantified in the peripheral blood of patients treated on clinical trial NCT03595917 via CAP/CLIA-approved clinical *BCR::ABL1* qRT-PCR performed in the clinical molecular laboratory of Brigham and Women’s Hospital (Boston, MA).

#### Targeted DNA Sequencing

PDX models underwent mutational profiling with targeted panels. Leukemia cells were enriched from fresh primary PDX bone marrow or peripheral blood via immunomagnetic enrichment for human B cells using human CD19 MicroBeads (Miltenyi Biotec #130-050-301; Gaithersburg, MD, USA). DNA was extracted using the DNeasy Blood & Tissue kit (QIAGEN #69504) and fluorometrically quantitated using the Qubit dsDNA HS assay kit (Invitrogen #Q32854; Waltham, MA, USA) prior to use in next-generation sequencing library preparation.

A hybrid-capture target enrichment panel targeting the full coding sequences of 183 genes selected based on the presence of recurrent mutations in hematologic malignancies was utilized to profile most PDX models at baseline, on-treatment, and at end of study (as previously described).^63^ An amplicon-based clinical sequencing panel targeting hotspot regions of the oncogenes and most of the coding regions of tumor suppressor genes recurrently implicated in hematologic malignancies (total 93 genes) was employed for a subset of PDX models.^64^ A custom amplicon-based deep sequencing panel targeting 23 genes implicated in in B-ALL treatment resistance (ArcherDX; Boulder, CO, USA) was employed to profile PDXs progressing after BCR::ABL1 inhibition.

#### Whole Exome Sequencing (WES) sample preparation

PDXs that progressed in absence of treatment-emergent driver alterations detected by targeted sequencing underwent whole exome sequencing using the SureSelect Human All Exon v5 kit (Agilent Life Sciences; Santa Clara, CA, USA). Briefly, 100ng of genomic DNA from each leukemia specimen as well as a control cell line (CEPH 1408) and a tail clipping from a non-xenografted NSG mouse were fragmented to 250bp on a Covaris Ultrasonicator (Woburn, MA, USA). Size-selected DNA fragments were ligated to xGen v1 UDI-UMI9 adaptors (Integrated DNA Technologies; Coralville, IA, USA) during automated library preparation with a Biomek FX^p^ liquid handling robot (Beckman Coulter; Indianapolis, IN, USA). Libraries (250ng per sample) were pooled to 750ng and captured with the SureSelect Human All Exon v5 bait set. Captures were pooled and sequenced on a HiSeq 3000 (Illumina; San Diego, CA, USA).

### Flow sorting of from healthy human bone marrow aspirates and PDX tumors

Approximately 10^6^ cells per sample were resuspended in PBS with 4,6-diamidino-2-phenylindole (DAPI; 0.75μg/mL) as a dead cell marker. For cell surface staining, PBS-washed cells were blocked with Fc blocker for 10 min on ice and then stained with the antibodies listed in **Table S10** at the manufacturers’ recommended concentrations or with an isotype control for 25 min on ice. Cells were then washed and resuspended in chilled PBS containing 0.75μg/mL of DAPI to exclude dead cells. For annexin V staining, annexin V binding buffer (BD Biosciences) was used instead of PBS, and 7-aminoactinomysin D (7-AAD; BD Biosciences) instead of DAPI. Phycoerythrin (PE)-labelled annexin V was purchased from BD Biosciences. Acquisition was performed on a LSR Fortessa flow cytometer (BD Biosciences). Fluorescence-based cell sorting was performed on a FACSAria II (BD Biosciences). FACS data were analyzed with FlowJo software (FlowJo).

Cells expressing B cell lineage-defining surface proteins were enriched by FACS on a BD FACSAria II cell sorter (BD Biosciences; Franklin Lakes, New Jersey, USA) based on staining with antibodies targeting the following markers: Annexin V, CD45, CD34, CD10, CD19, CD20, and CD22. Healthy and immunophenotyped subpopulations were defined as in **Figures S4A & S7B**. Lymphoid progenitor sub-populations then underwent scRNA-seq via Seq-Well S^3^ and SS2.

### Sample preparation for scRNA-seq of clinical and PDX samples

We used the Seq-Well S^3^ platform for massively parallel scRNA-seq to capture transcriptomes of single cells on barcoded mRNA capture beads.^31^ Briefly, a single-cell suspension of 15,000 cells in 200μL RPMI media supplemented with 10% FBS was loaded onto single arrays containing barcoded mRNA capture beads (ChemGenes). The arrays were sealed with a polycarbonate membrane (pore size of 0.01μm), before undergoing cell lysis and transcript hybridization. The barcoded mRNA capture beads were then recovered and pooled for all subsequent steps. Reverse transcription was performed using Maxima H Minus Reverse Transcriptase (Thermo Fisher Scientific EP0753). Exonuclease I treatment (NEB M0293 L) was used to remove excess primers, followed by Second Strand Synthesis using a primer of eight random bases to create complementary cDNA strands with SMART handles for PCR amplification. Whole transcriptome amplification was carried out using KAPA HiFi PCR Mastermix (Kapa Biosystems KK2602) with 2000 beads per 50-μl reaction volume. Libraries were then pooled in sets of eight (totaling 16,000 beads), purified using Agencourt AMPure XP beads (Beckman Coulter, A63881) by a 0.6× solid phase reversible immobilization (SPRI) followed by a 1× SPRI, and quantified using Qubit hsDNA Assay (Thermo Fisher Scientific Q32854). The quality of whole transcriptome amplification (WTA) product was assessed using the Agilent High Sensitivity D5000 Screen Tape System (Agilent Genomics) with an expected peak at 800 base pairs tailing off to beyond 3000 base pairs and a small/nonexistent primer peak.

Libraries were constructed using the Nextera XT DNA tagmentation method (Illumina FC-131–1096) on a total of 750pg of pooled cDNA library from 16,000 recovered beads using index primers with format as previously described.^31^ Tagmented and amplified sequences were purified at a 0.6× SPRI ratio yielding library sizes with an average distribution of 300 to 750bp in length as determined using the Agilent High Sensitivity D5000 Screen Tape System (Agilent Genomics). Two arrays were sequenced per sequencing run with an Illumina 75 Cycle NextSeq 500/550 v2 kit (Illumina FC-404–2005) at a final concentration of 2.4pM. The read structure was paired end with Read 1 starting from a custom Read 1 primer containing 20 bases with a 12-bp cell barcode and 8-bp unique molecular identifier (UMI) and Read 2 containing 50 bases of transcript sequence.

### Sample preparation for paired SMR mass profiling and SMART-Seq2

For all PDX and healthy bone marrow samples, cells were adjusted to a final concentration of 2.5*10^5^ cells/ml to load single cells into the mass sensor array and record single-cell mass measurements, as previously described.^41,65^ In order to exchange buffer and flush individual cells from the system, the release side of the device was constantly flushed with PBS at a rate of 15μL per minute. Upon detection of a single-cell at the final cantilever of the SMR, as indicated by a supra-threshold shift in resonant frequency, a set of three-dimensional motorized stages (ThorLabs) was triggered to move a custom PCR-tube strip mount from a waste collection position to a sample collection position to retrieve the cell. Each cell was dispensed in approximately 5μl of PBS into a PCR tube containing 5μl of 2× TCL lysis buffer (Qiagen) with 2% *v*/*v* 2-mercaptoethanol (Sigma) for a total final reaction volume of 10μl. After each 8-tube PCR strip was filled with cells, the strip was spun down at 1,000 g for 30 seconds and immediately snap-frozen on dry ice. Following collection, samples were stored at -80 C prior to library preparation and sequencing.

Single-cell lysates were compiled from independent collections upon thawing and transferred into wells of a 0.2mL skirted 96-well PCR plate (Thermo Fisher Scientific). scRNA-seq libraries were generated using SMART-Seq2 protocol.^66^ Briefly, cDNA was reversed transcribed from single cells using Maxima RT (Thermo Fisher Scientific) and whole transcriptome amplification (WTA) was performed. WTA products were purified using the Agencourt AMPure XP beads (Beckman Coulter) and used to prepare paired-end libraries with Nextera XT (Illumina). Single cells were pooled and sequenced on a NextSeq 550 sequencer (Illumina) using a 75 cycle High Output Kit (v2.5) with a 30bp paired end read structure.

## QUANTIFICATION AND STATISTICAL ANALYSIS

### PDX *in vivo* studies: survival analysis on treatment arms and with pretreatment clinical risk stratification metadata

Analyses fitting a Cox proportional hazards model for overall survival (OS) and progression-free survival (PFS) outcomes on treatment arms and pretreatment clinical risk stratification categories were performed using the *survival* package in R.^67^ The following pre-clinical features included: *IZKF1* deletion, 9p deletion, hyperdiploid karyotype, gain of chromosome 21, presenting white blood cell count, age, sex (if age <18 years), race, phase of disease, number of prior therapies, and pre-existing *ABL1* mutation(s). Hazard ratios and p-values for PFS within pretreatment clinical risk categories were generated relative to the lowest risk group in each category (**Figure S1D**).

### WES alignment and variant calling

Pooled sequenced WES samples were demultiplexed using Picard tools. Read pairs were aligned to the hg19 reference build using the Burrows-Wheeler Aligner.^68^ Data were sorted and duplicate-marked using Picard tools. Alignments were refined using the Genome Analysis Toolkit (GATK)^69,70^ for localized realignment around small insertion and deletion (indel) sites. Mutation analysis for single nucleotide variants was performed with MuTect v1.1.4^71^ and annotated by Variant Effect Predictor.^72^ Indels were called using the SomaticIndelDetector tool of the GATK. Copy number variants (CNVs) were identified using RobustCNV for autosomes.^73^ Detected alterations are reported in **Table S3** and **Figure S2A**.

### scRNA-seq sequencing alignment and quality control

Sequenced Seq-Well BCL files were demultiplexed into individual sample FASTQs for Read 1 and Read 2 using the bcl2fastq pipeline on Terra, as previously described. The resultant paired read FASTQs were aligned to the hg19 genome using the cumulus/dropseq_tools pipeline on Terra maintained by the Broad Institute using standard settings, generating a genes by cells count matrix for each sample.^74^ Low quality cells were filtered using nGene≤200, nUMI≤500, and percent mitochondrial transcripts≤30% thresholds before merging samples; genes were filtered if they were not expressed in at least 10 cells.

Sequenced SS2 BCL files were similarly demultiplexed using bcl2fastq and aligned to the hg19 genome using publicly available scripts on Terra (github.com/broadinstitute/TAG-public). Total gene counts and transcript per million (TPM) matrices were filtered to remove low quality cells with <15% transcriptome mapping, 2,000 genes, and 45,000 mapped reads, before continuing analysis. Genes expressed in fewer than 10 cells, as well as long non-coding RNAs and unique hg19 reference-build variants were removed before downstream analysis.

### Human healthy bone marrow reference cell type clustering and visualization

After QC filtering, 13,643 high quality cells from 7 healthy human bone marrow donors were analyzed in Seurat v2.3.4 to classify hematopoietic cell types.^75^ After normalization, the top 1,500 highly dispersed variable genes were selected using the mean-variance plot method in Seurat’s FindVariableFeatures function. ScRNA-seq data was scaled over highly variable genes and used as input for PCA analysis. The top significant PCs, as defined by the JackStraw test (top 25 PCs), were used as input for building a SNN graph to cluster cells by their (k=35) nearest neighbors and for t-SNE visualization of clusters. Given the shared, continuous hierarchy of covarying gene expression in hematopoietic development, broad cell types (progenitor, myeloid, erythroid, B cell lineage, pDCs, T cells, and Plasmablasts) were called based on their differentially expressed genes (identified using the Wilcox test in Seurat’s FindAllMarkers function), and subset into individual Seurat objects for a second round of clustering to resolve the final 13 cell types defined in **Figure S4**. Cell type annotations were *post-hoc* validated based on biased or exclusive expression of known marker genes (**Figure S4D**).

SS2 healthy reference cell types were called by their confident random forest prediction probabilities (see next section) and examination of marker genes to provide further support of cell type identification (**Figure S8B**). Cell type clusters were visualized using SPRING, a tool that generates force-directed layouts from kNN graphs to visually preserve hierarchical relationships between cell types.^76^

### Unbiased identification of consensus intratumoral gene expression programs with NMF

We sought to identify common axes of covarying intratumoral gene expression within all Ph+ ALL tumors in our dataset. First, we ran consensus NMF (cNMF) on each tumor in our dataset (n=52 total samples, defining bone marrow and spleen samples from the same mouse as individual tumors).^77^ For this analysis, we selected a consensus 1,489 variable genes across all tumors by first identifying the top 2,500 variable genes within each individual tumor using the variance standardized transformation method in Seurat v5.0.2 FindVariableFeatures function. To ensure consensus variable gene selection was not biased by PDX line- or patient-specific variable genes, as some models or donors had more tumors sampled than others, we initially selected the top 2,000 median weighted variable genes across tumors within a PDX line or patient, and then chose the top 2,000 median weighted variable genes across all PDX line and patient median gene lists. 511 of these top 2,000 variable genes were removed based on non-zero expression across all 52 tumors.

cNMF (1,000 iterations) was performed on the counts matrices of each tumor utilizing the consensus variable gene list over a range of k=3-9. All stable solutions of k, defined by a cNMF solution silhouette score>0.8 across iterations, were evaluated for optimal k selection using the following heuristics. We first hierarchically clustered the Jaccard Similarity of the top 50 genes from each factor across all stable k solutions; under-clustered k solutions were nominated based on factors that contained genes split across clusters that were hierarchically clustered in higher k factorizations, and over-clustered k solutions were nominated based on the presence of factors that did not hierarchically cluster with lower k factorizations or split genes across multiple lower k factors.^78^ To further evaluate these hypothesized over- or under-clustered k solutions, we scaled the data and ran UMAP projections over the top 50 genes from each factor for each stable k solutions. We used Seurat’s AddModuleScore function over the top 50 genes from each factor to assess whether under-clustered factors convolved expression across UMAP subclusters of optimal k solutions, or whether over-clustered factors scored highest in the same subcluster of cells or mostly strongly defined 1-2 cells (“junk” factor). Finally, we assessed significant Pearson correlation of the top 50 genes in each optimal k factor over an expression-binned bootstrapped null distribution as previously described,^79^ removing factors that were not significantly correlated (typically “activity”-like continuous programs in UMAP projections that, upon inspection, actually contained sparsely expressed genes of redundant biological annotations to other factors within that k solution). Factors from the selected optimal k that contained significantly correlated genes were labeled as “intratumoral gene expression programs” or GEPs, and collated for downstream intertumoral comparisons across the entire tumor cohort. Examples of intratumoral GEPs from representative PDX and patient tumors are shown in **Figure S3B**.

From performing intratumoral cNMF on 52 tumors, we identified 166 intratumoral GEPs. We excluded outlier GEPs by constructing a kNN graph (k=15) and filtered 40 intratumoral GEPs using an elbow-based filtering criterion over kNN distances of each individual GEP to its nearest neighbor. The remaining 126 intratumoral GEPs were hierarchically clustered using Ward.D clustering over their cosine similarity to reveal 7 meta-GEPs or “mGEPs”, which we interpret as shared intratumoral gene covariation across at least 8 individual tumors (**Figure S3A**). To interpret shared gene covariation across each identified mGEP, we isolated the top 30 median gene loadings across intratumoral GEPs within a given mGEP cluster (**Figures 3C & S3A; Table S4**).

### Training and interpreting the random forest classifier

Random forest is an ensemble machine learning method used for both classification and regression. Like other ensemble models, random forests combine multiple weak classifiers, in this case shallow decision trees, to make predictions. In this work, a random forest was used for classification. Here, we interrogate aberrant developmental hierarchies in ALL by using random forests to predict the nearest cell type from the normal B-cell lineage for single cells from Ph+ ALL samples. There are inherent advantages to random forests for the Ph+ ALL classification task. Importantly, ensemble classifiers, like a random forest, provide a distribution of class probabilities reflecting the similarity of each cell to each cell type the model was trained on. This is done by calculating the proportion of trees voting for a cell type for each given observation. To generate a single prediction for a cell, the highest-class probability becomes the prediction. The higher the probability of the chosen class, the more transcriptionally similar the cell is to that stage of B cell development. The distribution of class probabilities itself can be used to understand the certainty – or uncertainty – of a prediction. We leveraged this measure of uncertainty in predictions to evaluate how well a tumor cell fits a specific stage in B cell lineage (**Figure 2H**). A tumor cell with a more uniform distribution of probabilities over classes likely shares transcriptional features with many a wider range of stages of B cell development, potentially indicating a more aberrant cell from normal development. Second, ensemble approaches tend to be more robust to overfitting, which is necessary when applying a model trained on sorted, healthy populations of cells to evaluate aberrant leukemic cells. Finally, because random forests are nonparametric models, they also are highly flexible to input feature scale and variance. This makes the approach particularly suited to raw count matrices output by various scRNA-seq technologies used.

Here, we trained a random forest on sorted cells from the B cell lineage using 15,000 genes with detected expression in more than 10 cells as input features. Random forests were implemented using R version 3.5.1 using the caret package for training infrastructure.^80^ The ranger implementation of random forests was used.^81^ Hyperparameter search over ranger parameters (the number of randomly selected features considered for splitting at each tree node and the rule used for splitting) was done via 10-fold cross-validation (CV). The model achieved an accuracy of 94±0.006% on 10-fold CV with optimal parameters. The final model used the full training set of 13,643 cells. Results of 10-fold cross validation are provided in **Figure S5A**. The model was also evaluated on an external testing set of Seq-Well generated healthy bone marrow scRNA-seq transcriptomes,^16^ and achieved performance of average AUC=0.99 over all 13 cell types (**Figure S5C**). To interpret features being used to make predictions by the classifier, we used permutation importance tests. Permutation importance measures the impact of randomly shuffling feature values on the performance of a model measured as accuracy and decrease in Gini impurity. Specifically, a computationally accelerated heuristic method was used that constructs a null distribution from features that have importance values close to zero, limiting the need for randomly shuffling all features independently to evaluate significance.^82^ The results of feature importance defining marker genes segregating the 13 cell types can be found in **Figure S5B**.

### Generating Tumor Hybrid Scores and assigning leukemia cells to hybrid populations

Tumor Hybrid gene signatures were generated as previously described.^16^ First, normalized gene expression values were correlated to RF cell type classification probabilities along B cell progenitor cell types (HSC, Pre-B, and Immature B). Pro-B RF probability correlations were excluded; since most leukemic cells were dominantly classified as Pro-B with secondary classifications along B cell lineage cell types, genes that highly correlated to Pro-B RF probabilities were not Pro-B-specific. To ensure that genes in each hybrid population signature were specific and unique to HSC, Pre-B, and Immature B cell types, the second-highest cell type correlation coefficient was subtracted from the highest correlation coefficient for a given cell type. Additionally, to ensure that cell type signatures were not obfuscated by cell cycle, positive correlation values of genes with cell cycle scores were subtracted from the highest correlation coefficient of a given cell type. After performing these corrections, the top 30 correlated genes to HSC, Pre-B, and Immature B cell types were included in their respective hybrid gene signatures; a threshold of 30 genes was selected based on the approximate elbow in corrected correlation values for each hybrid signature. Likewise, Pro-B gene scores were defined by the top 30 differentially expressed genes in healthy Pro-B cells (**Figures S6A; Table S5**).

Tumor cells were scored by these HSC, Pro-B, Pre-B, and Immature B gene signatures using the Seurat v4 AddModuleScore function, and consequently assigned to hybrid populations similarly to what has been described previously.^60^ Single cells were classified into HSC-like, PreB-like, and Immature-like hybrid populations based on their highest hybrid cell type signature score, which we required to be > 0.5 + that cell’s Pro-B score. All other cells were classified as Pro-B like cells, which were characterized by strong Pro-B gene expression and weak or no co-expression of other cell type hybrid signature scores. The classifications based on these hybrid score distributions and relative to their B cell lineage RF prediction probabilities is demonstrated in **Figure S6B**.

### Mutual information of transcription factor activities with tumor hybrids

We sought to elucidate gene programs whose activity associated with the tumor hybrid populations defined above. Given the highly entropic co-expression of tumor hybrid signatures with Pro-B marker genes, we utilized mutual information as a metric for the potentially non-linear mutual dependence of gene expression with hybrid-defined developmental marker genes. Within respective hybrid subpopulations of each individual PDX line’s pre-treatment and progression time points, we calculated the average normalized mutual information (NMI) of all highly expressed genes across the top 30 genes in each hybrid population signature, using raw gene counts as input. Within each PDX sample and hybrid population, MI values between each gene-gene pair were generated using R infotheo package mutinformation function with the Miller-Madow asymptotic bias corrected empirical estimator and normalized to scale values between 0 and 1 as a relative, comparable metric between samples.^83^ We interpret these NMI values as a metric for genes whose expression relatively scale with hybrid population identity.

To identify cooperatively expressed genes that are collectively mutually informed with tumor hybrid signatures, we utilized the collectRI transcription factor accessibility database along with the decoupleR package to *in silico* predict mutually informed transcription factor (TF) activity with tumor hybrid identity.^84^ Averaged NMI values for each PDX sample hybrid were used as input with the run_ulm function to estimate the linear relationship between TF-target genes and their hybrid marker gene expression. Within each PDX samples, significant TFs were ordered by their variance in mutually informed activity between hybrid populations, and the top 30 of these TFs were selected for further inspection of scaled predicted activity between hybrid subsets. NMI values and *in silico* predicted TF activities for each healthy reference population (HSC for HSC-hyb, Pre-BI and Pre-BII for PreB-hyb, Immature B for ImmB-hyb) were generated analogously and *post-hoc* compared to their leukemic hybrid counterparts (subset shown in **Figure S6D**), demonstrating that the majority of leukemic hybrid-defining TF activities were conserved with their healthy counterparts, with a couple of TFs.

### Defining developmental skews in Smart-seq2 PDX samples

Given the paucity of RF-classified immature B cells in the SS2 leukemic dataset (**Figure S8A**), we identified genes that were Pearson correlated with Pre-B RF probabilities and with progenitor population (HSC, GMP, Pro-Mono, Early-Erythroid) probabilities. We found that genes correlated with progenitor RF probabilities negatively correlated with Pre-B RF probabilities in leukemic cells and vice versa, enabling us to define a spectrum of differentiation between progenitor and later-stage B cell developmental stages (**Figures S8C & S8D**). Progenitor-like and PreB-like scores were generated by scoring leukemic cells over the top 30 genes significantly correlated to their respective RF probabilities (**Table S7**). Each cell’s location on the leukemic differentiation spectrum was defined by its (PreB-like score – Progenitor-like score).

### Identifying somatic variants in full-length Smart-seq2 (SS2) scRNA-seq libraries

Each sample’s SS2 FASTQ files were aligned to hg19 using STAR (version 2.6.0c) and then sorted and indexed with SAMtools (version 1.13).^85,86^ 16 genomic loci, nominated based on recurrently identified SNVs from bulk RNA-seq in the genes *KRAS, NRAS, PTPN11, GNB1, ABL1,* and *STAT5A* (**Figure S9A; Table S3**), were assessed for wild-type or mutant transcript detection by a custom script utilizing the Pysam library (version 0.16.0.1).^87^ In particular, for each locus of interest, each cell was marked as “NC” if there was no coverage at the locus, marked with 0 if all overlapping reads matched the reference allele, or marked as mutant if there were overlapping reads that did not match the reference allele.

### Predicting chromosomal number variations (CNVs) in SS2 scRNA-seq libraries with inferCNV

To identify SS2 leukemic cells harboring CNVs and *in silico* elucidate subclonal heterogeneity within tumors, we estimated single-cell CNVs as previously described by computing the average expression in a sliding window of 100 genes within each chromosome after sorting the detected genes by their hg19 genome-defined chromosomal coordinates.^88,89^ We used all healthy bone marrow SS2 cells identified above (**Figure S8B**) as reference normal populations for this analysis. Complete information on the inferCNV workflow used for this analysis can be found here: https://github.com/broadinstitute/inferCNV/wiki, using baseline input parameters for SS2 data and for the i6 HMM algorithm for confident CNV-positive or negative predictions in single-cells.

### Module scoring single-cell transcriptomes

Module scores of all gene signatures over single-cells were annotated using the Seurat v4 AddModuleScore function, which calculates the average expression levels of genes in a gene list relative to all other genes with comparable normalized gene expression. Quiescent cells were binned based on positive scores for a literature-derived quiescence gene signature derived from human hematopoietic cells.^90^ We utilized previously established signatures for G1/S (n=43 genes) and G2/M (n=55 genes) to place each cell along this dynamic process;^89^ after inspecting the distribution of scores in the complete dataset, we considered any cell > 1.5 SD above the mean for either the G1/S or the G2/M scores to be cycling.^16^ Senescence scores were derived from the top 50 genes significantly differentially expressed in the SS2 DFAB-25157 RAS-mutant cells in remission compared to all other RAS-mutant SS2 cells (**Figure 4F; Table S8**).

### Defining stress-autophagy, pre-BCR signaling, and inflammation transcriptional programs at remission

To define heterogeneous, correlated transcriptional states defining PDX tumors that emerge in MRD, we first performed differential gene expression analysis between paired pre-treatment and MRD cells within the same PDX line to identify genes that significantly increase expression at remission. A total of 40 MRD state-defining genes were identified based on significant upregulation in at least two PDX-specific MRD differentially expressed gene (DEG) lists. Performing gene-gene Pearson correlation across the expression of these 40 shared MRD-high DEGs in all remission leukemic cells revealed three correlated modules of genes. To expand these three modules, we identified the top 30 genes significantly correlated (>2 standard deviations above median Pearson correlation) with the top differentially expressed gene in each module (**Table S9**). Pathway enrichment of significantly correlated genes was performed over msigDB Reactome gene sets for functional annotation, and to nominate targeted inhibitors of state (**Figures S10C**).

## SUPPLEMENTARY INFORMATION

**Table S1. Clinical characteristics of patients whose tumors were used to generate PDX models.**

*Related to Figure 1*

**Table S2. Characteristics of PDX models.**

*Related to Figure 1*

**Table S3. Detected Alterations in PDX Leukemias.**

*Related to Figure 1*

**Table S4. cNMF meta-GEP gene lists.**

*Related to Figure 2*

**Table S5. Seq-Well derived Tumor Hybrid signatures.**

*Related to Figure 2*

**Table S6. Patient characteristics and clinical trial outcomes**.

*Related to Figure 3*

**Table S7. SS2-derived Tumor Hybrid signatures.**

*Related to Figure 4*

**Table S8. Senescence-Like signature.**

*Related to Figures 4*

**Table S9. MRD State signatures.**

*Related to Figure 5*

**Table S10. Flow cytometry antibodies.**

*Related to Methods*

**Figures S1-11.**

---

**Figure S1.**
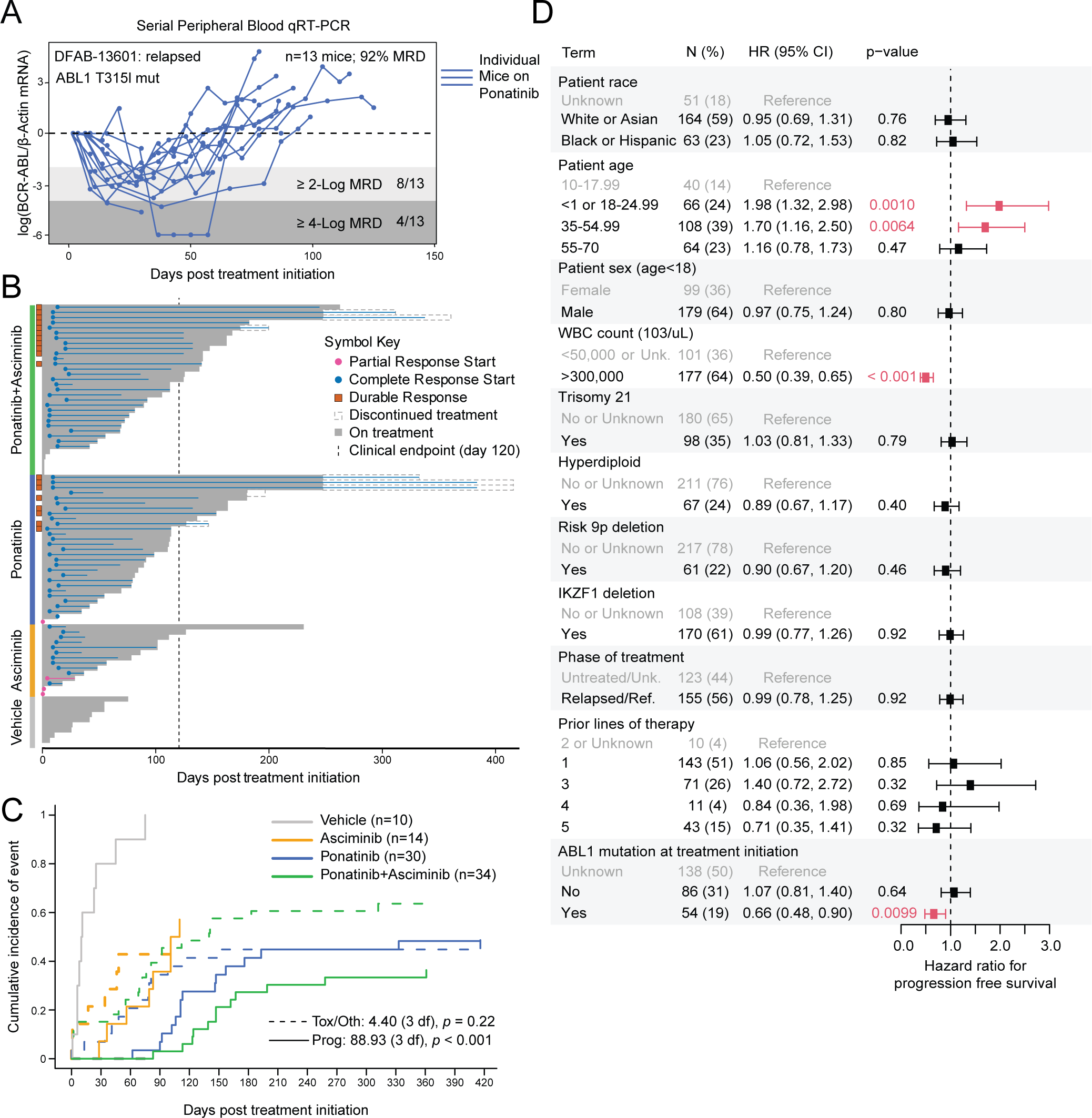
*In vivo* PDX Phase II-like trial outcomes. *Related to Figure 1* **(A)** Serial peripheral blood *BCR::ABL1* qRT-PCR measurements from PDX model DFAB-13601 treated with Ponatinib daily (40mg/kg/day); each line represents an individual mouse. **(B)** Key trial events and outcomes for each mouse on Phase II-like trial, grouped by treatment arm. Complete response indicates <4% peripheral blood circulating blasts detected via flow cytometry; partial response indicates reduced peripheral blood blasts compared to pretreatment but >1% involvement; durable response indicates complete remission past 120 days on therapy. **(C)** Competing risks model comparing progression and non-progression mortality in mice by treatment arm; p-values from a Cox regression analysis indicated for differences in progression and non-progression outcomes between treatment arms. **(D)** Hazard ratios comparing pre-clinical risk factors for progression free survival in PDX mice (see **Methods**). Significant shifts (*p*<0.05 from Cox regression analysis) annotated in red. Median hazard ratios plotted with error bars representing ±1 quartile; “N”=number of mice; “HR”=hazard ratio; “CI”=confidence interval.

**Figure S2.**
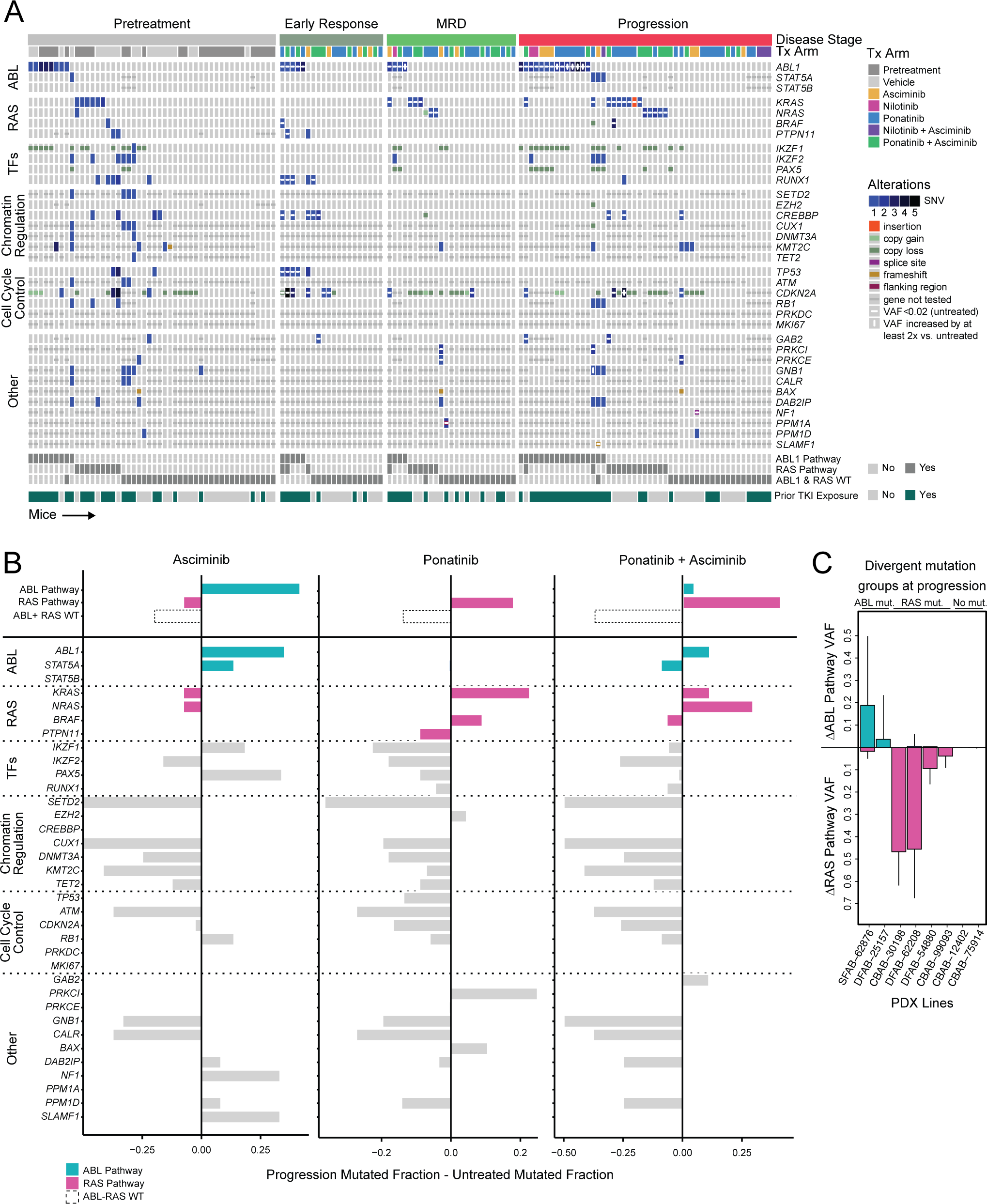
Emergent patterns in *BCR::ABL1* B-ALL mutation acquisition on TKI. *Related to Figure 1* **(A)** Mutational alterations of individual PDX mice on TKI therapeutic regimen, grouped by disease stage and annotated by treatment arm (“Tx Arm”). Treatment emergent mutations indicated when mice from the same PDX line were profiled at pretreatment. Summary of grouped RAS or ABL pathway mutations included below. Mice are annotated for prior TKI exposure. “MRD”=minimal residual disease; “TFs”=transcription factors. Alteration details additionally reported in **Table S3**. **(B)** Change in the fraction of mice on each Phase II-like trial treatment arm that harbor mutations between progression and pretreatment. Genes along the ABL and RAS pathways are annotated in turquoise and magenta, respectively. **(C)** Change in average VAF of PDX lines at progression for mutations along the ABL or RAS pathways compared to paired pretreatment tumors. Error bars indicate +1 standard deviation from the plotted mean ΔVAF.

**Figure S3.**
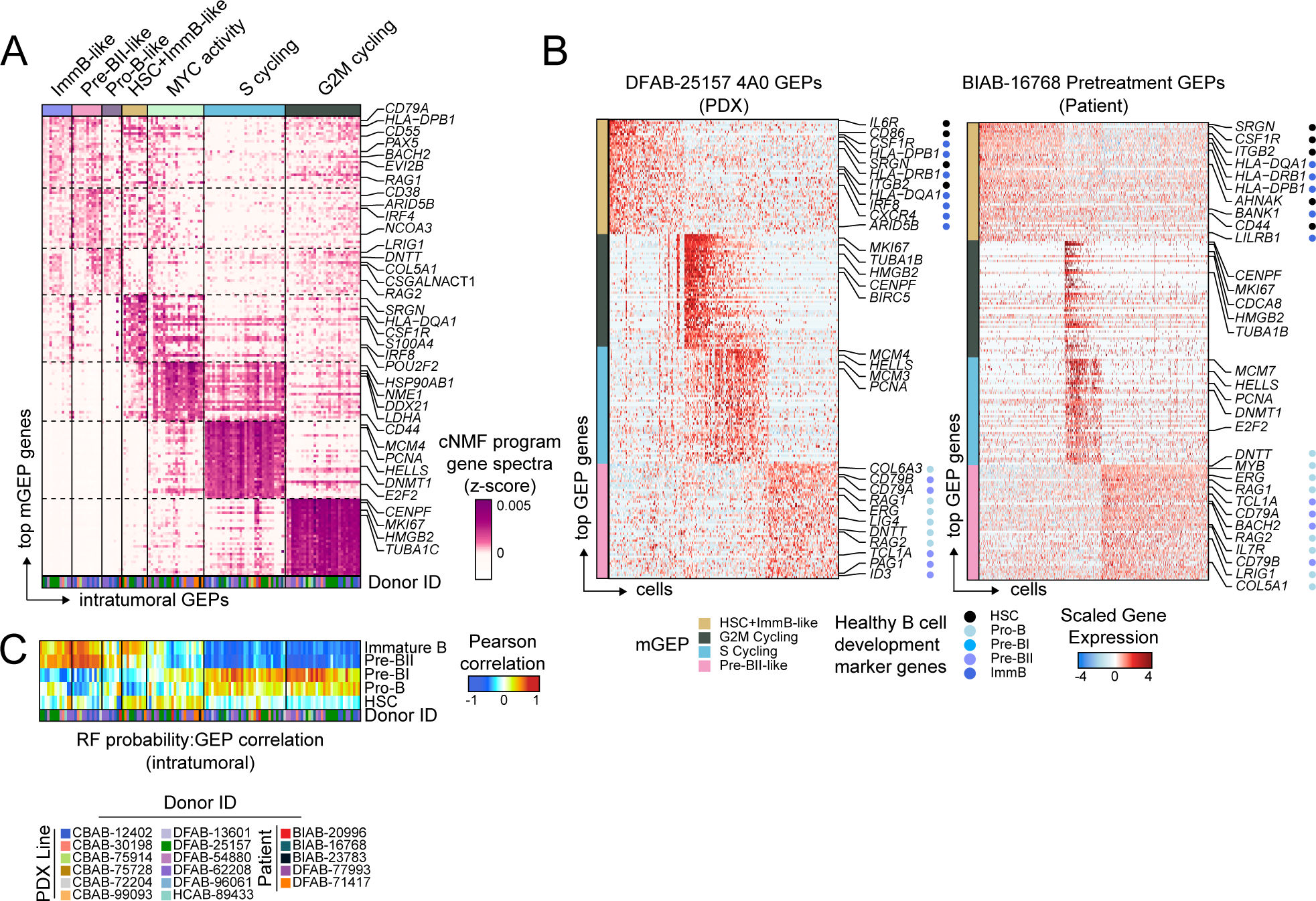
Intratumoral cNMF reveals developmentally convolved gene co-expression. *Related to Figure 2* **(A)** cNMF program z-scored gene spectra for the top 30 metaprogram (mGEP) genes across all intratumoral gene expression programs (GEPs; Table S4); individual GEPs are annotated by PDX or Patient ID to show mGEP consensus across multiple donors. **(B)** Representative heatmaps demonstrating intratumoral GEPs for one PDX tumor (DFAB-25157 4A0) and one patient tumor (BIAB-16768 Pretreatment). Known, healthy B cell lineage marker genes are annotated for each GEP. **(C)** Pearson correlation of GEP module score and random forest (RF) classification probabilities. Bottom color track indicates the donor where each individual GEP was identified.

**Figure S4.**
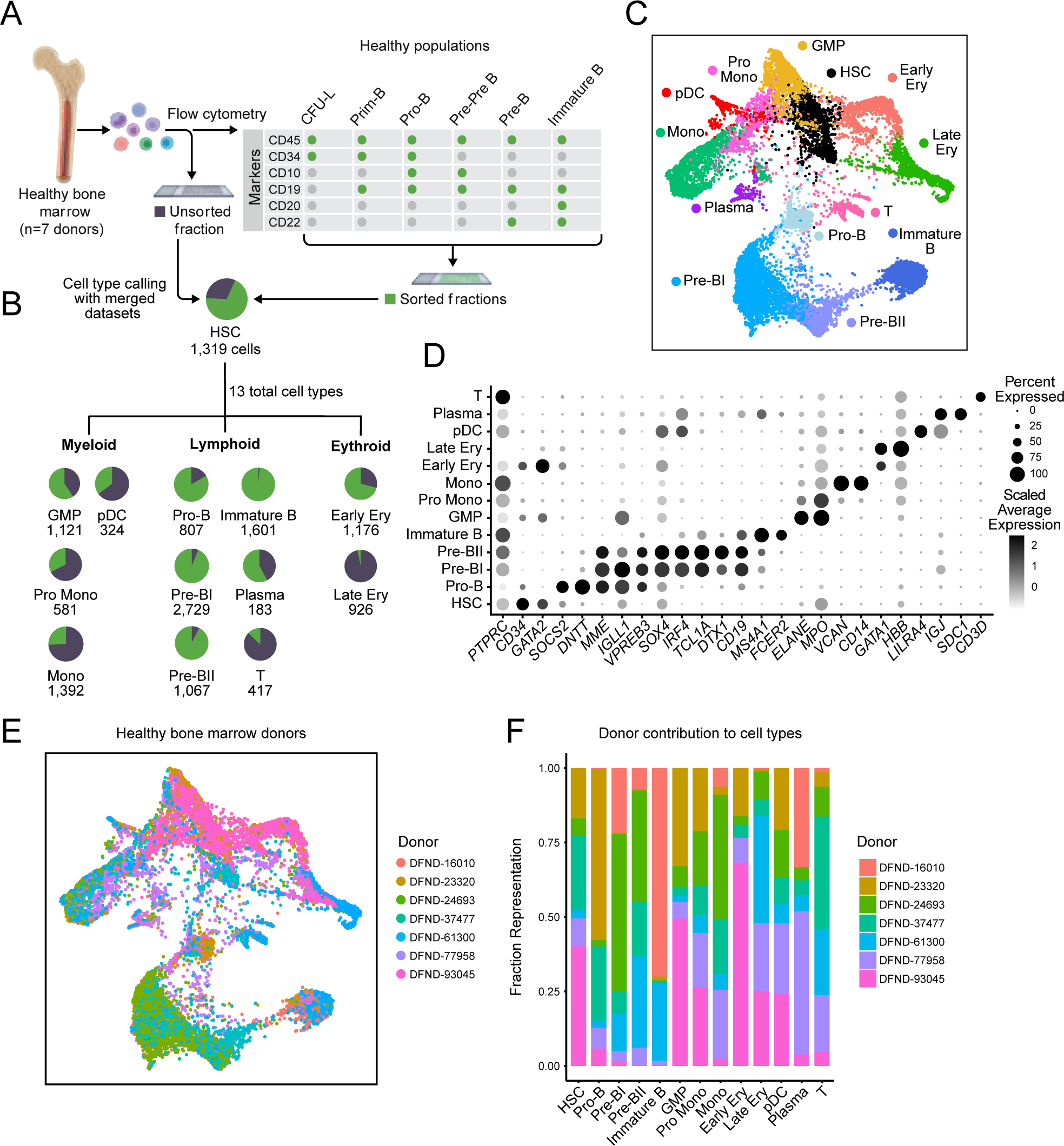
Generation of healthy human bone marrow scRNA-seq dataset. *Related to Figure 2* **(A)** Healthy human bone marrow samples (n = 7) were flow sorted into live bulk, CFU-L (colony-forming unit low; progenitor), Prim-B, Pro-B, Pre-Pre-B, Pre-B, and Immature B populations for scRNA-seq profiling (see Methods). **(B)** Proportion of each cell type identified from the bulk (gray) or flow sorted-fraction (green). **(C)** Force-directed graph (FDG) projection of healthy human bone marrow annotated by hematopoietic cell types (n=13,643 cells). **(D)** Dot plot of hematopoietic cell type marker genes. Color denotes scaled average expression; size denotes percent expression in each scRNA-seq cell type population. **(E)** FDG projection of healthy human bone marrow, annotated by donor. **(F)** Donor fractional contribution to each cell type population.

**Figure S5.**
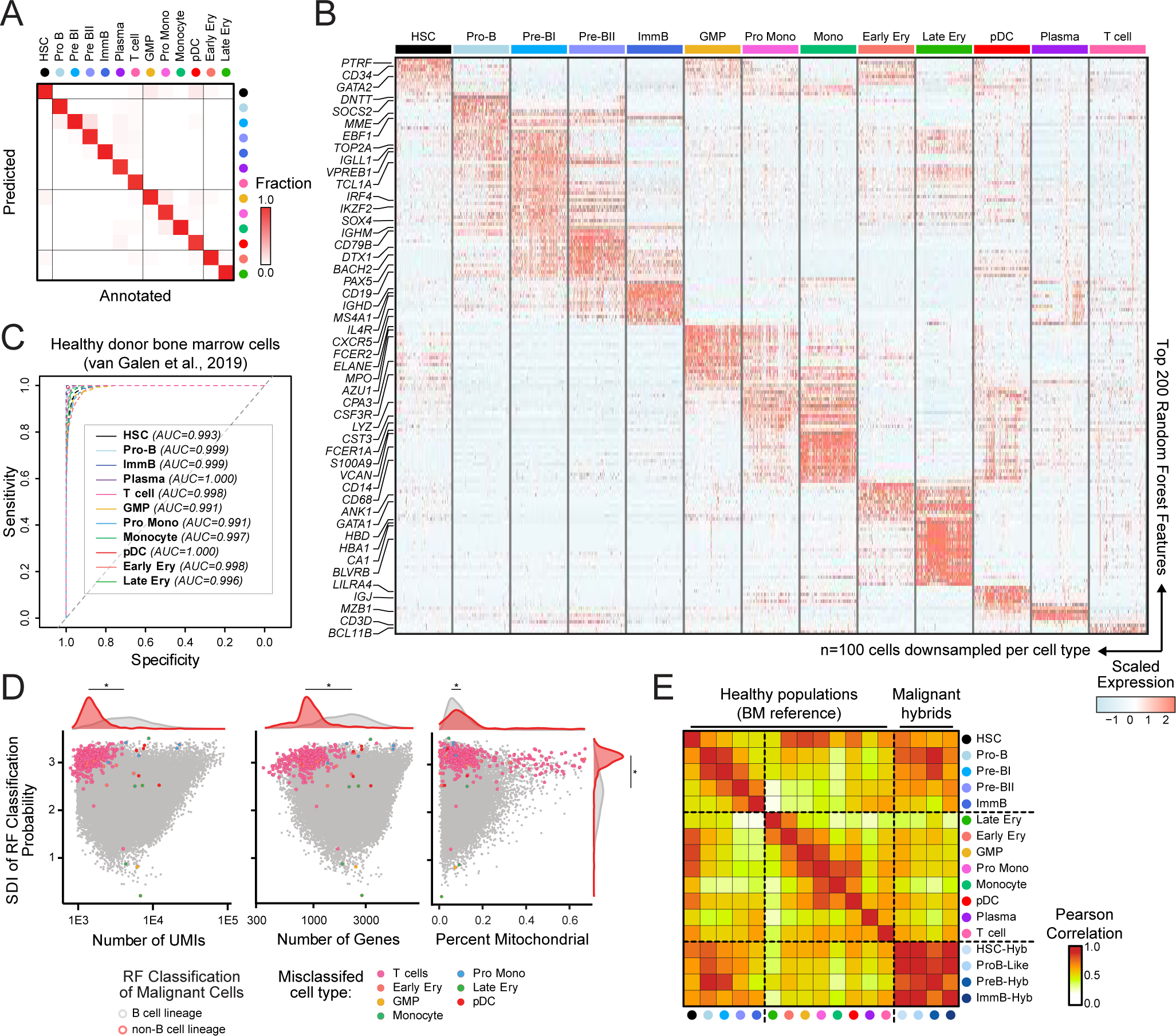
Random Forest Classifier accurately classifies healthy and Ph+ ALL single-cell transcriptomes. *Related to Figure 2* **(A)** 10-fold cross-validation of each healthy reference cell type during RF training. **(B)** Top 200 RF features ranked by permuted feature importance, grouped by healthy reference cell type (randomly down-sampled n=100 cells). **(C)** Receiver Operating Characteristics (ROC) curves for RF classification of test scRNA-seq bone marrow dataset;^16^ area under the ROC curve (AUC) values listed in inset for each cell type. **(D)** Shannon Diversity Index (SDI) of classification probabilities versus number of unique molecular identifier (UMI), number of genes, and percent mitochondrial transcripts for all leukemic cells. Cells removed from analysis due to highest non-B cell lineage classification are outlined in red and colored by misclassified cell type. Significant shifts in distribution between non-B lineage and B-lineage single-cells, as defined by a Kolmogorov-Smirnov (KS) test, reported (\**p*<0.001). **(E)** Pearson correlation over gene expression of top 2,000 highly-variable genes from healthy reference dataset across healthy and malignant hybrid cell type subpopulations.

**Figure S6.**
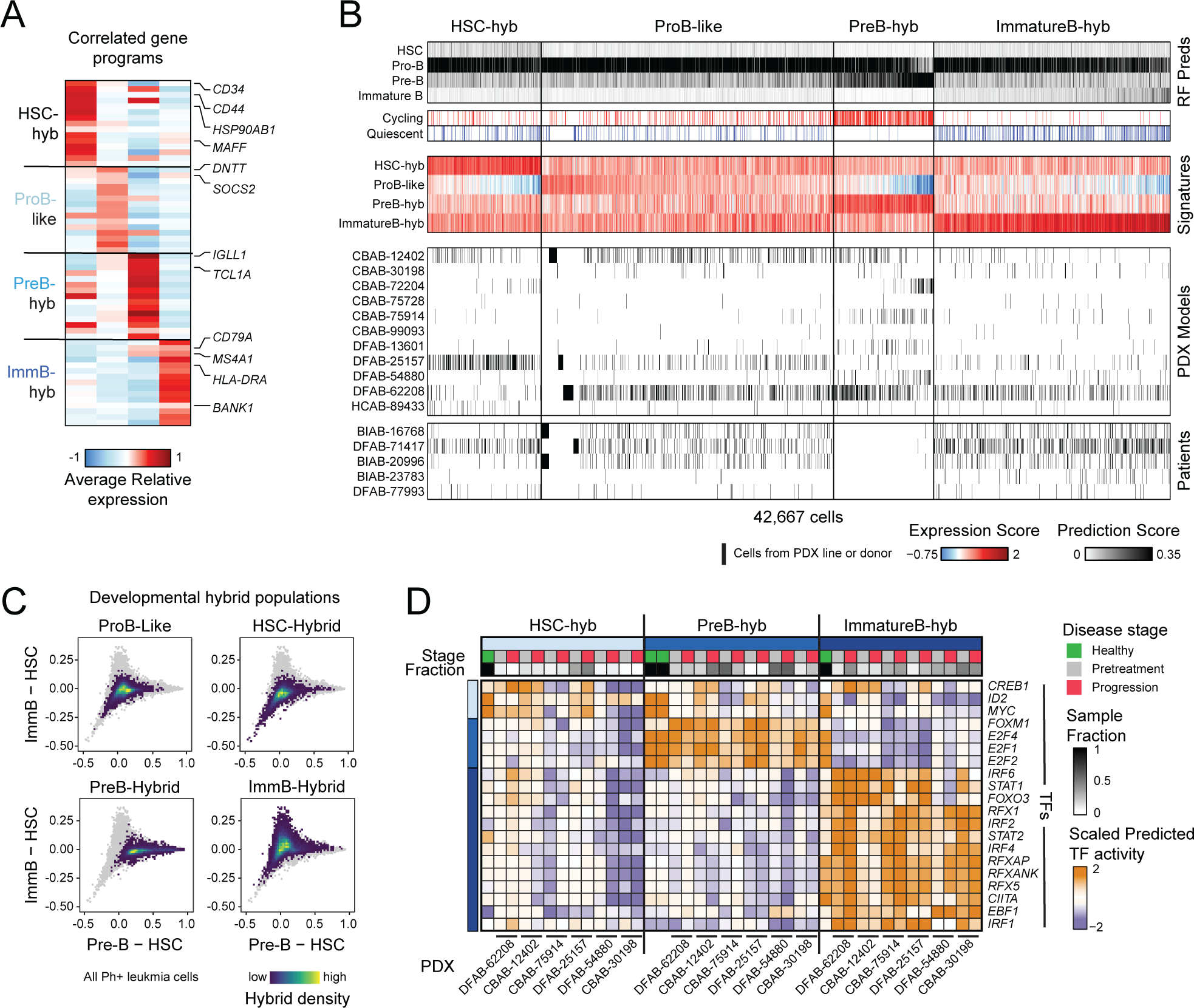
Defining Ph+ ALL developmental tumor hybrid populations. *Related to Figure 2* **(A)** Developmental hybrid signatures defined by the top 30 genes correlated to RF prediction scores for each normal B-lineage cell type (Table S5). Average expression of signature genes across leukemic hybrid populations. **(B)** Classification of leukemic hybrid populations based on random forest (RF) classification probabilities and hybrid signatures (see Methods). RF prediction probabilities, cycling or quiescent status, and PDX line or Patient ID annotated for each cell. **(C)** Leukemic hybrid subpopulations projected onto RF prediction probability axes, as in **Figure 2H**. Densities of leukemia cells from each hybrid population projected over the landscape of all leukemia cells in the scRNA-seq dataset (plotted in grey). **(D)** Scaled *in silico* predicted transcription factor (TF) activity over genes associated with developmental hybrid gene signatures (see **Methods**). Scaled TF activity scores shown in human reference samples (green) and PDX lines at pretreatment (grey) and progression (red), subset to TFs whose predicted activity scale with HSC, Pre-B, and Immature B RF classification probabilities in leukemic cells. Healthy reference Pre-BI and Pre-BII populations plotted independently within Pre-B.

**Figure S7.**
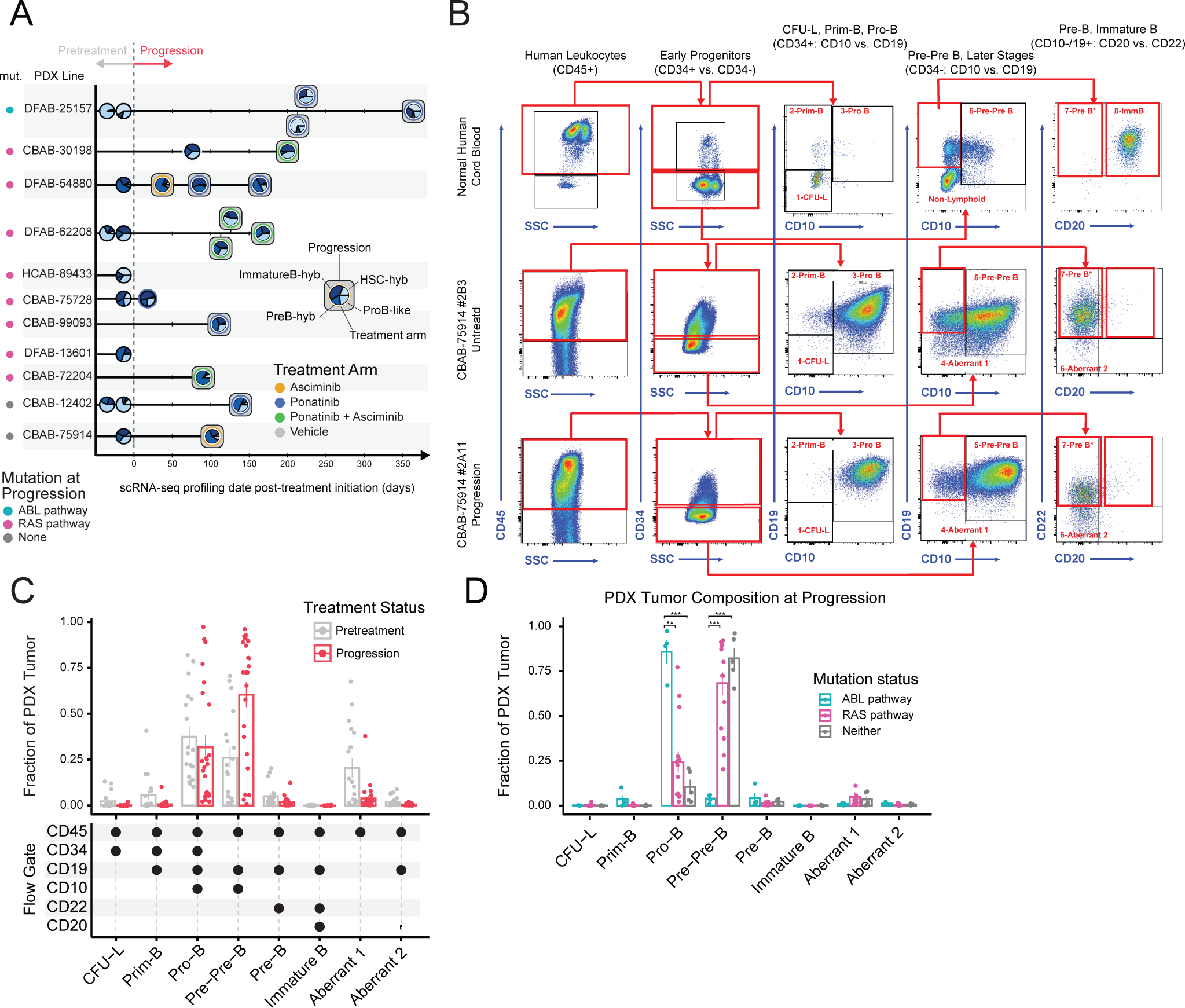
Transcriptional and immunophenotype shifts on therapy. *Related to Figure 3* **(A)** Hybrid scRNA-seq population distributions for each profiled pretreatment and progression PDX mouse, annotated by treatment arm and time on treatment. **(B)** Flow sorting gating strategy for B cell progenitor populations on a representative healthy human umbilical cord blood sample, PDX pre-treatment tumor, and PDX progression tumor (representative PDX=CBAB-75914). **(C)** Fraction representation of PDX pretreatment and progression tumors across immunophenotyped B cell progenitor-like populations. Individual tumor immunophenotyped population fractions plotted as points; bars represent average tumor fraction within each immunophenotyped population at pretreatment or progression time points, including error bars for ±1 standard deviation. Surface markers used for flow gating of each population, as shown in (B), annotated below. **(D)** Fraction of PDX tumor at progression of each immunophenotyped B cell progenitor-like population, grouped by mutation status at progression; bars represent average tumor fraction, with error bars for ±1 standard deviation. Significant p-values from Dirichlet regression noted; \*\**p*<0.01 and \*\*\**p*<0.001.

**Figure S8.**
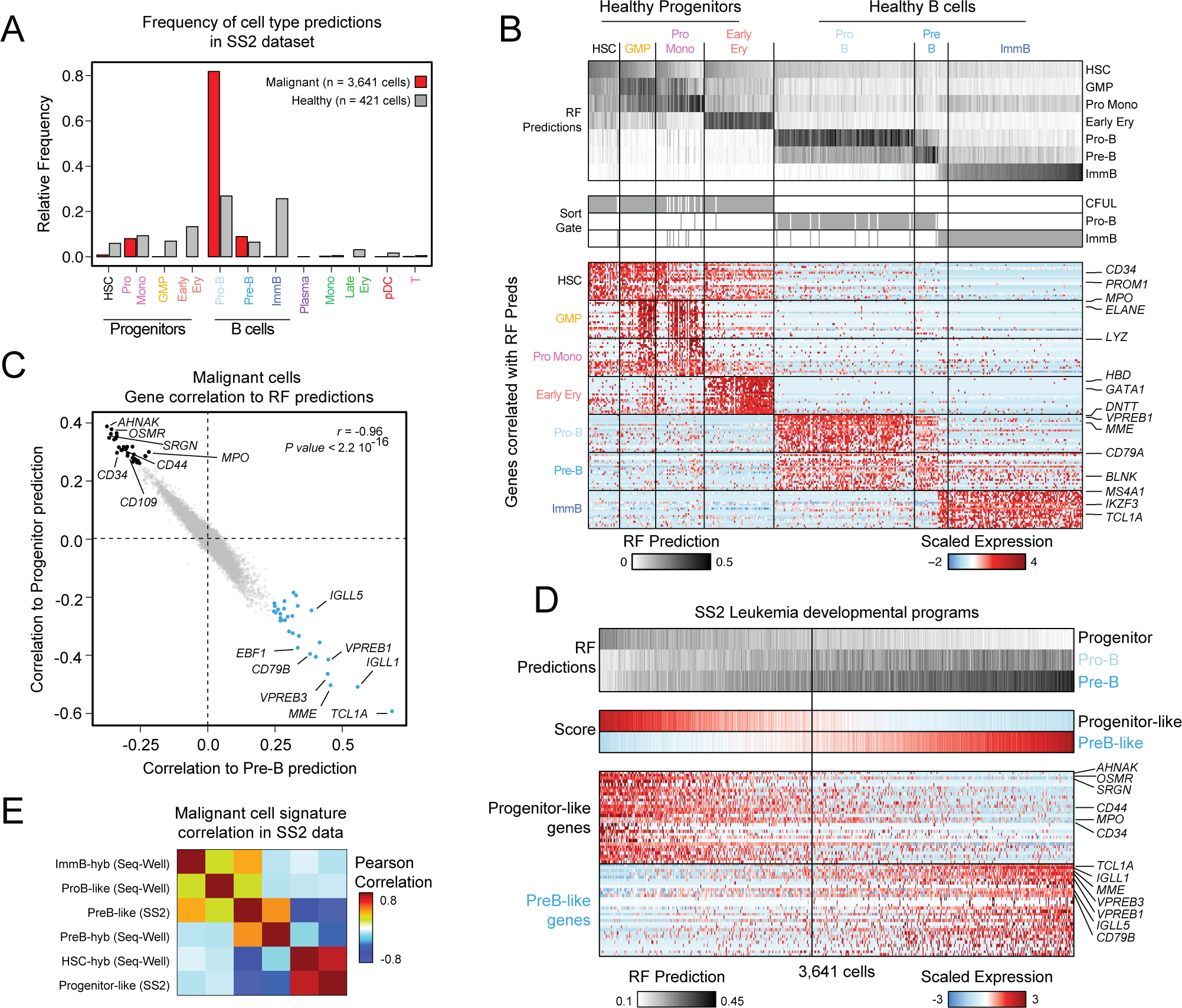
Random Forest (RF) Classifier recovers developmental structure in Smart-Seq2 single-cell transcriptomes. *Related to Figure 4* **(A)** Proportion of RF cell type classifications across all Smart-Seq2 (SS2) healthy and leukemic cells. **(B)** Single-cells ordered by RF prediction probabilities from progenitor cell types to differentiated B cell types, and annotated by flow sort gate (as in Figure S4A). Below, scaled expression of the top 10 RF prediction-correlated genes in developmentally-ordered healthy cells. **(C)** Genes correlated to Pre-B RF prediction (x-axis) and genes correlated to Progenitor RF prediction are negatively correlated with each other; rho and p-value from Pearson correlation noted. Colored points represent the top 30 progenitor and Pre-B correlated genes used to define the SS2 developmental spectrum. **(D)** Leukemic SS2 single-cells ranked by Progenitor-like score, annotated by B cell lineage RF prediction probabilities. Below, scaled expression of top 30 Progenitor-like and PreB-like signature genes (Table S7). **(E)** Pearson cross-correlation of RF cell type-correlated gene signature scores derived from Seq-Well and SS2 show cross-modality concordance. For clarity, SS2 signatures are hereafter referenced as “HSC-hyb” for Progenitor-like scores, and “PreB-hyb” for PreB-like scores.

**Figure S9.**
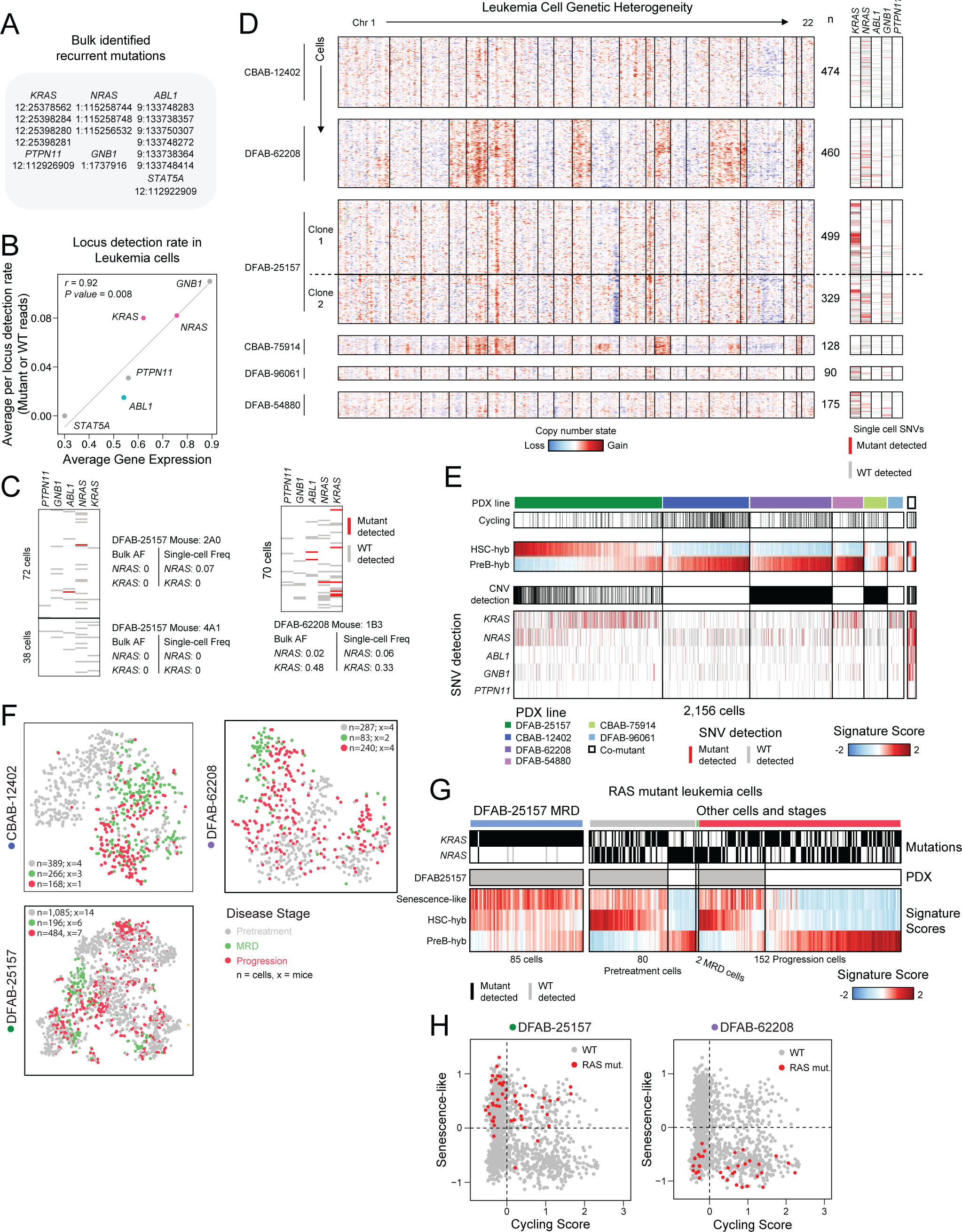
SS2 enables co-detection of mutations and transcriptome in leukemic single-cells. *Related to Figure 4* **(A)** Summary of recurrently-identified RAS-pathway and ABL-pathway mutation loci from bulk targeted sequencing across PDX lines that were aligned for mutation detection in SS2 FASTQs (see Methods; Table S3). **(B)** For genes with recurrently-identified mutations, Pearson correlation of average gene expression and normalized mutation-locus detection rate (either mutant or wild-type reads). **(C)** Mutant and wild-type transcripts detected in SS2 single-cell transcriptomes from three representative PDX tumors; detected mutant transcript frequency in single-cells matched bulk VAF. **(D)** Single-cell CNV profiles across each PDX line, including instances of CNV subclonal heterogeneity, paired with SS2-detected SNVs. **(E)** SS2 single-cells within each profiled PDX line ordered by HSC-hyb expression scores, as defined in Figure S8D. Cycling status, CNV detection, and detected mutant and wild-type transcripts are annotated. Co-mutant indicates single-cells where RAS and ABL pathway mutations were detected. **(F)** t-SNE projection of SS2 single-cells from representative PDX lines CBAB-12402, DFAB-62208, and DFAB-25157, colored by treatment time point. Number of SS2-profiled cells and mice at each time point denoted (n=cells, x=mice). **(G)** All RAS-pathway mutant leukemic single-cells grouped by three treatment timepoints, annotated by *KRAS* or *NRAS* mutant transcript detection, and ordered by HSC-hyb signature scores within SS2 single-cells from DFAB-25157 and non-DFAB-25157 PDX lines demonstrates association between senescence-like and HSC-hyb gene expression scores across PDX lines and treatment stages. **(H)** Cells from DFAB-25157 and DFAB-62208 at MRD and Progression, plot along fitness quadrants as defined in Figure 4H, with RAS-mutant leukemia cells annotated in red.

**Figure S10.**
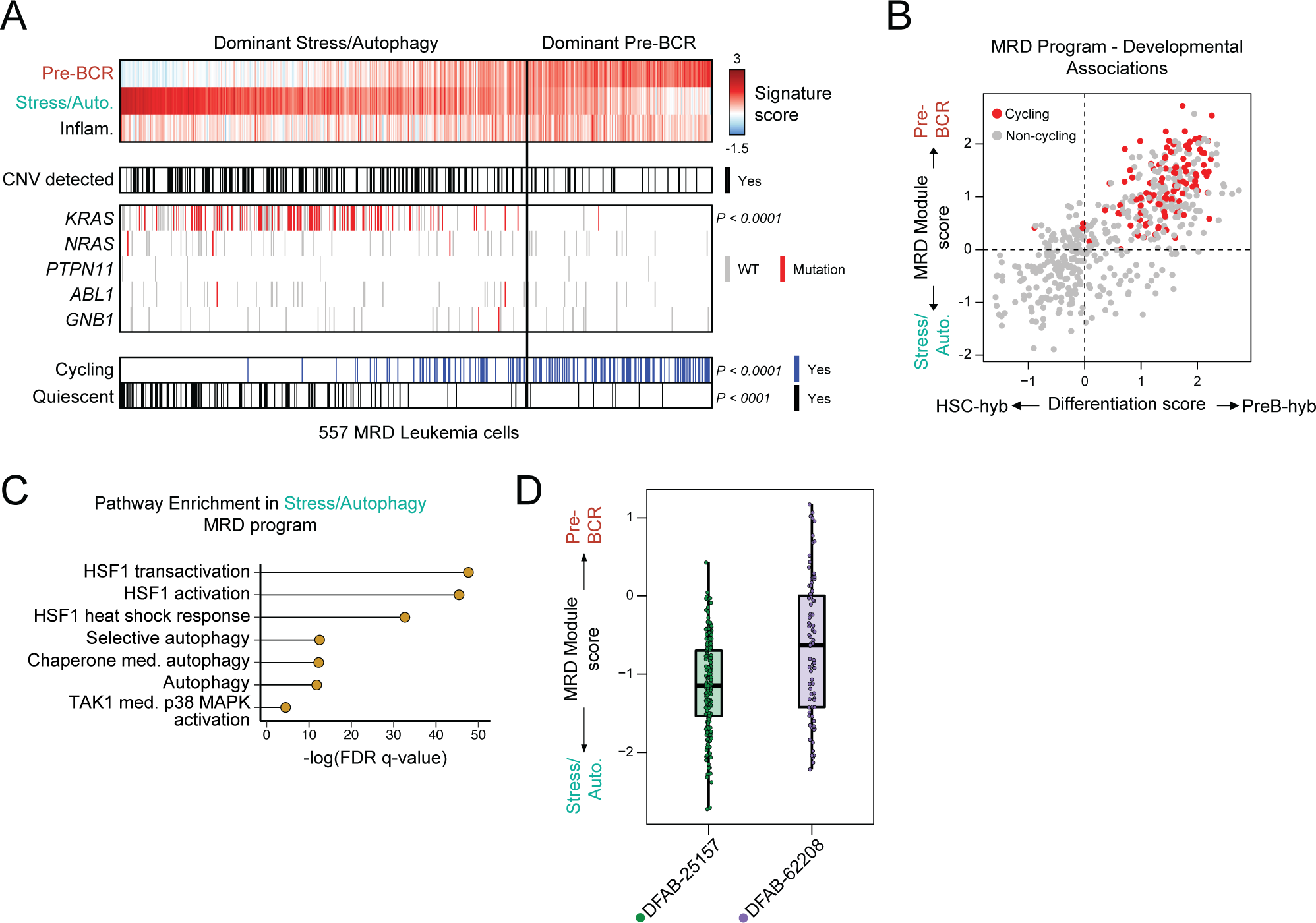
Targeting integrative cell states enhances remission. *Related to Figure 5* **(A)** All MRD single-cells ordered by Pre-BCR Signaling MRD state scores. CNV and SNV mutation status annotated for each cell, along with cycling and quiescent status. *P*-values reported from Fisher exact test comparing abundance of *KRAS*-mutant, quiescent, and/or cycling MRD cells with dominant Stress/Autophagy (“Stress/Auto.”) expression scores to those with dominant Pre-BCR Signaling expression scores. **(B)** Correlation between Stress/Autophagy and HSC-hyb gene expression, versus Pre-BCR signaling and PreB-hyb gene expression. Cycling cells annotated in red. **(C)** Pathway enrichment false discovery rate (FDR) q-values for the top 100 genes in the Stress/Autophagy MRD state. **(D)** Boxplot of relative MRD program (Pre-BCR Signaling – Stress/Autophagy) in MRD cells from DFAB-25157 and DFAB-62208; single-cell scores from each PDX-line plotted as individual points.

**Figure S11.**
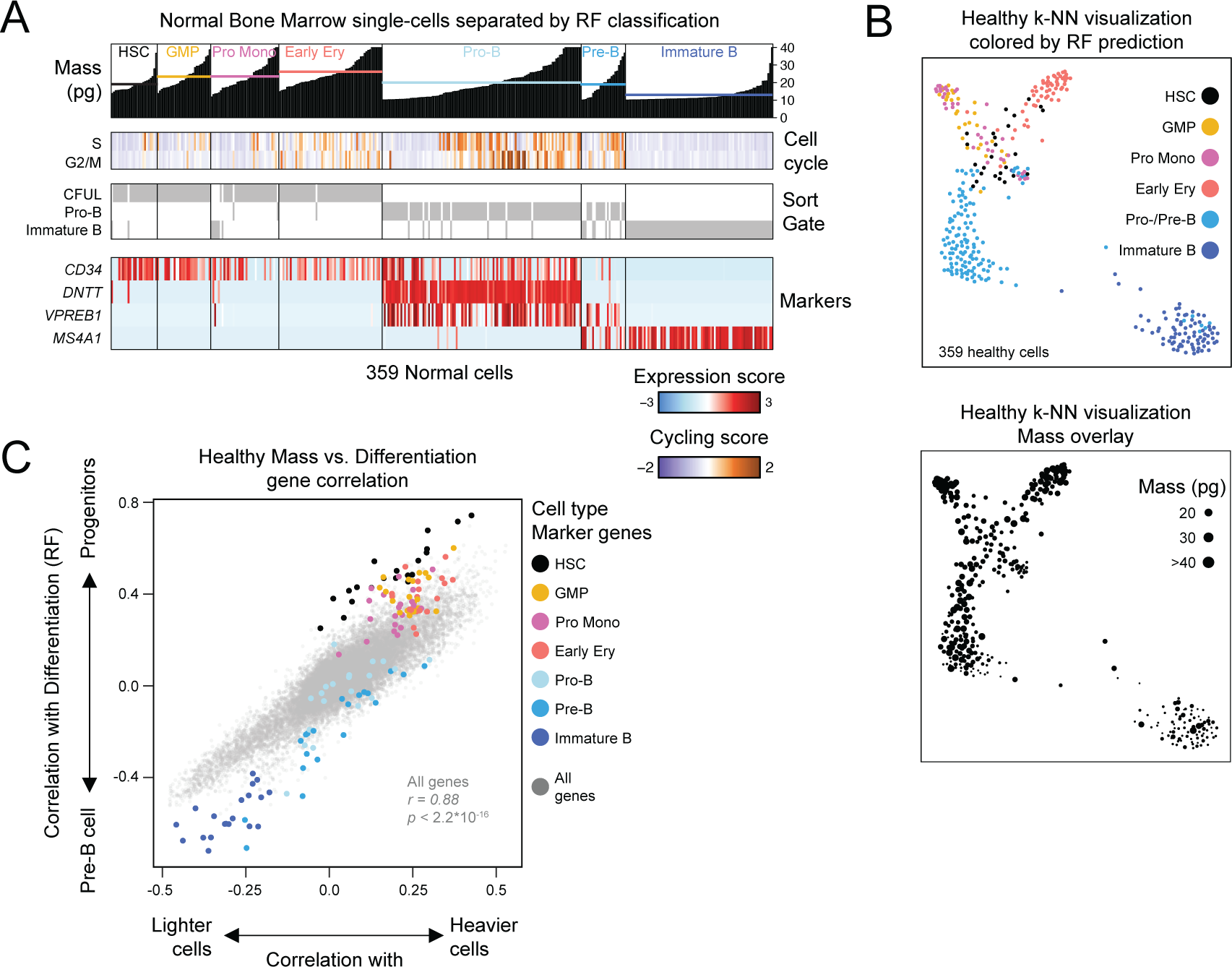
Mass correlates with developmental states and cell cycle. *Related to Figure 6* **(B)** Mass of healthy reference SS2 cells, binned by random forest-classified cell type and annotated by cell-type marker gene expression. Mean mass for each cell type plotted as a line. **(C)** Force directed graph (FDG) visualization of healthy SS2 cells, annotated by cell type (top) and by cell mass (bottom); dot size indicates cell mass. **(D)** Mass-correlated genes in healthy SS2 cells on the x-axis, versus the difference between genes correlated with RF progenitor and Pre-B cell types in healthy SS2 cells on the y-axis. Colored points denote marker genes for each cell type. R and p-value denote Pearson correlation between x- and y-axis indicated gene correlations.

## REFERENCES

1. Byrgazov, K., Lucini, C.B., Valent, P., Hantschel, O., and Lion, T. (2018). BCR-ABL1 compound mutants display differential and dose-dependent responses to ponatinib. Haematologica 103, e10–e12. 10.3324/haematol.2017.176347.

2. Soverini, S., De Benedittis, C., Papayannidis, C., Paolini, S., Venturi, C., Iacobucci, I., Luppi, M., Bresciani, P., Salvucci, M., Russo, D., et al. (2014). Drug resistance and BCR-ABL kinase domain mutations in Philadelphia chromosome–positive acute lymphoblastic leukemia from the imatinib to the second-generation tyrosine kinase inhibitor era: The main changes are in the type of mutations, but not in the frequency of mutation involvement. Cancer 120, 1002– 1009. 10.1002/cncr.28522.

3. Eide, C.A., Zabriskie, M.S., Savage Stevens, S.L., Antelope, O., Vellore, N.A., Than, H., Schultz, A.R., Clair, P., Bowler, A.D., Pomicter, A.D., et al. (2019). Combining the Allosteric Inhibitor Asciminib with Ponatinib Suppresses Emergence of and Restores Efficacy against Highly Resistant BCR-ABL1 Mutants. Cancer Cell 36, 431–443.e5. 10.1016/j.ccell.2019.08.004.

4. Bhang, H.C., Ruddy, D.A., Krishnamurthy Radhakrishna, V., Caushi, J.X., Zhao, R., Hims, M.M., Singh, A.P., Kao, I., Rakiec, D., Shaw, P., et al. (2015). Studying clonal dynamics in response to cancer therapy using high-complexity barcoding. Nat Med 21, 440–448. 10.1038/nm.3841.

5. Hochhaus, A., Kreil, S., Corbin, A.S., La Rosée, P., Müller, M.C., Lahaye, T., Hanfstein, B., Schoch, C., Cross, N.C.P., Berger, U., et al. (2002). Molecular and chromosomal mechanisms of resistance to imatinib (STI571) therapy. Leukemia 16, 2190–2196. 10.1038/sj.leu.2402741.

6. Good, Z., Sarno, J., Jager, A., Samusik, N., Aghaeepour, N., Simonds, E.F., White, L., Lacayo, N.J., Fantl, W.J., Fazio, G., et al. (2018). Single-cell developmental classification of B cell precursor acute lymphoblastic leukemia at diagnosis reveals predictors of relapse. Nat Med 24, 474–483. 10.1038/nm.4505.

7. Huang, X., Li, Y., Zhang, J., Yan, L., Zhao, H., Ding, L., Bhatara, S., Yang, X., Yoshimura, S., Yang, W., et al. (2024). Single-cell systems pharmacology identifies development-driven drug response and combination therapy in B cell acute lymphoblastic leukemia. Cancer Cell 42, 552–567.e6. 10.1016/j.ccell.2024.03.003.

8. Kim, J.C., Chan-Seng-Yue, M., Ge, S., Zeng, A.G.X., Ng, K., Gan, O.I., Garcia-Prat, L., Flores-Figueroa, E., Woo, T., Zhang, A.X.W., et al. (2023). Transcriptomic classes of BCR-ABL1 lymphoblastic leukemia. Nat Genet 55, 1186–1197. 10.1038/s41588-023-01429-4.

9. Bastian, L., Beder, T., Barz, M.J., Bendig, S., Bartsch, L., Walter, W., Wolgast, N., Brändl, B., Rohrandt, C., Hansen, B.-T., et al. (2024). Developmental trajectories and cooperating genomic events define molecular subtypes of BCR::ABL1-positive ALL. Blood 143, 1391– 1398. 10.1182/blood.2023021752.

10. Chan, L.N., Murakami, M.A., Robinson, M.E., Caeser, R., Sadras, T., Lee, J., Cosgun, K.N., Kume, K., Khairnar, V., Xiao, G., et al. (2020). Signalling input from divergent pathways subverts B cell transformation. Nature 583, 845–851. 10.1038/s41586-020-2513-4.

11. Ebinger, S., Özdemir, E.Z., Ziegenhain, C., Tiedt, S., Castro Alves, C., Grunert, M., Dworzak, M., Lutz, C., Turati, V.A., Enver, T., et al. (2016). Characterization of Rare, Dormant, and Therapy-Resistant Cells in Acute Lymphoblastic Leukemia. Cancer Cell 30, 849–862. 10.1016/j.ccell.2016.11.002.

12. Hurtz, C., Wertheim, G.B., Loftus, J.P., Blumenthal, D., Lehman, A., Li, Y., Bagashev, A., Manning, B., Cummins, K.D., Burkhardt, J.K., et al. (2020). Oncogene-independent BCR-like signaling adaptation confers drug resistance in Ph-like ALL. Journal of Clinical Investigation 130, 3637–3653. 10.1172/JCI134424.

13. Giustacchini, A., Thongjuea, S., Barkas, N., Woll, P.S., Povinelli, B.J., Booth, C.A.G., Sopp, P., Norfo, R., Rodriguez-Meira, A., Ashley, N., et al. (2017). Single-cell transcriptomics uncovers distinct molecular signatures of stem cells in chronic myeloid leukemia. Nat Med 23, 692–702. 10.1038/nm.4336.

14. Eadie, L.N., Saunders, V.A., Branford, S., White, D.L., and Hughes, T.P. (2018). The new allosteric inhibitor asciminib is susceptible to resistance mediated by ABCB1 and ABCG2 overexpression in vitro. Oncotarget 9, 13423–13437. 10.18632/oncotarget.24393.

15. Fennell, K.A., Vassiliadis, D., Lam, E.Y.N., Martelotto, L.G., Balic, J.J., Hollizeck, S., Weber, T.S., Semple, T., Wang, Q., Miles, D.C., et al. (2022). Non-genetic determinants of malignant clonal fitness at single-cell resolution. Nature 601, 125–131. 10.1038/s41586-021-04206-7.

16. van Galen, P., Hovestadt, V., Wadsworth II, M.H., Hughes, T.K., Griffin, G.K., Battaglia, S., Verga, J.A., Stephansky, J., Pastika, T.J., Lombardi Story, J., et al. (2019). Single-Cell RNA-Seq Reveals AML Hierarchies Relevant to Disease Progression and Immunity. Cell 176, 1265–1281.e24. 10.1016/j.cell.2019.01.031.

17. Velten, L., Story, B.A., Hernández-Malmierca, P., Raffel, S., Leonce, D.R., Milbank, J., Paulsen, M., Demir, A., Szu-Tu, C., Frömel, R., et al. (2021). Identification of leukemic and pre-leukemic stem cells by clonal tracking from single-cell transcriptomics. Nat Commun 12, 1366. 10.1038/s41467-021-21650-1.

18. Nam, A.S., Kim, K.-T., Chaligne, R., Izzo, F., Ang, C., Taylor, J., Myers, R.M., Abu-Zeinah, G., Brand, R., Omans, N.D., et al. (2019). Somatic mutations and cell identity linked by Genotyping of Transcriptomes. Nature 571, 355–360. 10.1038/s41586-019-1367-0.

19. Li, Q., Bohin, N., Wen, T., Ng, V., Magee, J., Chen, S.-C., Shannon, K., and Morrison, S.J. (2013). Oncogenic Nras has bimodal effects on stem cells that sustainably increase competitiveness. Nature 504, 143–147. 10.1038/nature12830.

20. Robinson, T.M., Bowman, R.L., Persaud, S., Liu, Y., Neigenfind, R., Gao, Q., Zhang, J., Sun, X., Miles, L.A., Cai, S.F., et al. (2023). Single-cell genotypic and phenotypic analysis of measurable residual disease in acute myeloid leukemia. Sci Adv 9, eadg0488. 10.1126/sciadv.adg0488.

21. Luskin, M.R., Murakami, M.A., Manalis, S.R., and Weinstock, D.M. (2018). Targeting minimal residual disease: a path to cure? Nat Rev Cancer 18, 255–263. 10.1038/nrc.2017.125.

22. Tettero, J.M., Ngai, L.L., Bachas, C., Breems, D.A., Fischer, T., Gjertsen, B.T., Gradowska, P., Griskevicius, L., Janssen, J.J.W.M., Juliusson, G., et al. (2023). Measurable residual disease-guided therapy in intermediate-risk acute myeloid leukemia patients is a valuable strategy in reducing allogeneic transplantation without negatively affecting survival. Haematologica 108, 2794–2798. 10.3324/haematol.2022.282639.

23. Dekker, S.E., Rea, D., Cayuela, J.-M., Arnhardt, I., Leonard, J., and Heuser, M. (2023). Using Measurable Residual Disease to Optimize Management of AML, ALL, and Chronic Myeloid Leukemia. Am Soc Clin Oncol Educ Book 43, e390010. 10.1200/EDBK_390010.

24. Wieduwilt, M.J. (2022). Ph+ ALL in 2022: is there an optimal approach? Hematology Am Soc Hematol Educ Program 2022, 206–212. 10.1182/hematology.2022000338.

25. Agudo, J., Aguirre-Ghiso, J.A., Bhatia, M., Chodosh, L.A., Correia, A.L., and Klein, C.A. (2024). Targeting cancer cell dormancy. Nat Rev Cancer 24, 97–104. 10.1038/s41568-023-00642-x.

26. Wylie, A.A., Schoepfer, J., Jahnke, W., Cowan-Jacob, S.W., Loo, A., Furet, P., Marzinzik, A.L., Pelle, X., Donovan, J., Zhu, W., et al. (2017). The allosteric inhibitor ABL001 enables dual targeting of BCR–ABL1. Nature 543, 733–737. 10.1038/nature21702.

27. Townsend, E.C., Murakami, M.A., Christodoulou, A., Christie, A.L., Köster, J., DeSouza, T.A., Morgan, E.A., Kallgren, S.P., Liu, H., Wu, S.-C., et al. (2016). The Public Repository of Xenografts Enables Discovery and Randomized Phase II-like Trials in Mice. Cancer Cell 29, 574–586. 10.1016/j.ccell.2016.03.008.

28. Gökbuget, N., Stanze, D., Beck, J., Diedrich, H., Horst, H.-A., Hüttmann, A., Kobbe, G., Kreuzer, K.-A., Leimer, L., Reichle, A., et al. (2012). Outcome of relapsed adult lymphoblastic leukemia depends on response to salvage chemotherapy, prognostic factors, and performance of stem cell transplantation. Blood 120, 2032–2041. 10.1182/blood-2011-12-399287.

29. Martinelli, G., Iacobucci, I., Storlazzi, C.T., Vignetti, M., Paoloni, F., Cilloni, D., Soverini, S., Vitale, A., Chiaretti, S., Cimino, G., et al. (2009). IKZF1 (Ikaros) deletions in BCR-ABL1-positive acute lymphoblastic leukemia are associated with short disease-free survival and high rate of cumulative incidence of relapse: a GIMEMA AL WP report. J Clin Oncol 27, 5202– 5207. 10.1200/JCO.2008.21.6408.

30. Pfeifer, H., Wassmann, B., Pavlova, A., Wunderle, L., Oldenburg, J., Binckebanck, A., Lange, T., Hochhaus, A., Wystub, S., Brück, P., et al. (2007). Kinase domain mutations of BCR-ABL frequently precede imatinib-based therapy and give rise to relapse in patients with de novo Philadelphia-positive acute lymphoblastic leukemia (Ph+ ALL). Blood 110, 727–734. 10.1182/blood-2006-11-052373.

31. Hughes, T.K., Wadsworth, M.H., Gierahn, T.M., Do, T., Weiss, D., Andrade, P.R., Ma, F., de Andrade Silva, B.J., Shao, S., Tsoi, L.C., et al. (2020). Second-Strand Synthesis-Based Massively Parallel scRNA-Seq Reveals Cellular States and Molecular Features of Human Inflammatory Skin Pathologies. Immunity 53, 878–894.e7. 10.1016/j.immuni.2020.09.015.

32. Baccin, C., Al-Sabah, J., Velten, L., Helbling, P.M., Grünschläger, F., Hernández-Malmierca, P., Nombela-Arrieta, C., Steinmetz, L.M., Trumpp, A., and Haas, S. (2020). Combined single-cell and spatial transcriptomics reveal the molecular, cellular and spatial bone marrow niche organization. Nat Cell Biol 22, 38–48. 10.1038/s41556-019-0439-6.

33. Slobodnyuk, K., Radic, N., Ivanova, S., Llado, A., Trempolec, N., Zorzano, A., and Nebreda, A.R. (2019). Autophagy-induced senescence is regulated by p38α signaling. Cell Death Dis 10, 376. 10.1038/s41419-019-1607-0.

34. Naka, K., Jomen, Y., Ishihara, K., Kim, J., Ishimoto, T., Bae, E.-J., Mohney, R.P., Stirdivant, S.M., Oshima, H., Oshima, M., et al. (2015). Dipeptide species regulate p38MAPK-Smad3 signalling to maintain chronic myelogenous leukaemia stem cells. Nat Commun 6, 8039. 10.1038/ncomms9039.

35. Sui, X., Kong, N., Ye, L., Han, W., Zhou, J., Zhang, Q., He, C., and Pan, H. (2014). p38 and JNK MAPK pathways control the balance of apoptosis and autophagy in response to chemotherapeutic agents. Cancer Lett 344, 174–179. 10.1016/j.canlet.2013.11.019.

36. Naeim, F., Song, S.X., Rao, P.N., and Phan, R.T. (2018). Atlas of Hematopathology: Morphology, Immunophenotype, Cytogenetics, and Molecular Approaches 2nd ed. (Elsevier, Inc).

37. Burg, T.P., Godin, M., Knudsen, S.M., Shen, W., Carlson, G., Foster, J.S., Babcock, K., and Manalis, S.R. (2007). Weighing of biomolecules, single cells and single nanoparticles in fluid. Nature 446, 1066–1069. 10.1038/nature05741.

38. Feijó Delgado, F., Cermak, N., Hecht, V.C., Son, S., Li, Y., Knudsen, S.M., Olcum, S., Higgins, J.M., Chen, J., Grover, W.H., et al. (2013). Intracellular water exchange for measuring the dry mass, water mass and changes in chemical composition of living cells. PLoS One 8, e67590. 10.1371/journal.pone.0067590.

39. Hecht, V.C., Sullivan, L.B., Kimmerling, R.J., Kim, D.-H., Hosios, A.M., Stockslager, M.A., Stevens, M.M., Kang, J.H., Wirtz, D., Vander Heiden, M.G., et al. (2016). Biophysical changes reduce energetic demand in growth factor-deprived lymphocytes. J Cell Biol 212, 439–447. 10.1083/jcb.201506118.

40. Cermak, N., Olcum, S., Delgado, F.F., Wasserman, S.C., Payer, K.R., A Murakami, M., Knudsen, S.M., Kimmerling, R.J., Stevens, M.M., Kikuchi, Y., et al. (2016). High-throughput measurement of single-cell growth rates using serial microfluidic mass sensor arrays. Nat Biotechnol 34, 1052–1059. 10.1038/nbt.3666.

41. Kimmerling, R.J., Prakadan, S.M., Gupta, A.J., Calistri, N.L., Stevens, M.M., Olcum, S., Cermak, N., Drake, R.S., Pelton, K., De Smet, F., et al. (2018). Linking single-cell measurements of mass, growth rate, and gene expression. Genome Biology 19, 207. 10.1186/s13059-018-1576-0.

42. Neurohr, G.E., Terry, R.L., Lengefeld, J., Bonney, M., Brittingham, G.P., Moretto, F., Miettinen, T.P., Vaites, L.P., Soares, L.M., Paulo, J.A., et al. (2019). Excessive Cell Growth Causes Cytoplasm Dilution And Contributes to Senescence. Cell 176, 1083–1097.e18. 10.1016/j.cell.2019.01.018.

43. Son, S., Tzur, A., Weng, Y., Jorgensen, P., Kim, J., Kirschner, M.W., and Manalis, S.R. (2012). Direct observation of mammalian cell growth and size regulation. Nat Methods 9, 910– 912. 10.1038/nmeth.2133.

44. Miettinen, T.P., Kang, J.H., Yang, L.F., and Manalis, S.R. (2019). Mammalian cell growth dynamics in mitosis. Elife 8, e44700. 10.7554/eLife.44700.

45. Marquart, J., Chen, E.Y., and Prasad, V. (2018). Estimation of the Percentage of US Patients With Cancer Who Benefit From Genome-Driven Oncology. JAMA Oncol 4, 1093–1098. 10.1001/jamaoncol.2018.1660.

46. Letai, A. (2017). Functional precision cancer medicine—moving beyond pure genomics. Nat Med 23, 1028–1035. 10.1038/nm.4389.

47. Letai, A., Bhola, P., and Welm, A.L. (2022). Functional precision oncology: Testing tumors with drugs to identify vulnerabilities and novel combinations. Cancer Cell 40, 26–35. 10.1016/j.ccell.2021.12.004.

48. Braun, T.P., Eide, C.A., and Druker, B.J. (2020). Response and Resistance to BCR-ABL1-Targeted Therapies. Cancer Cell 37, 530–542. 10.1016/j.ccell.2020.03.006.

49. Boumahdi, S., and de Sauvage, F.J. (2020). The great escape: tumour cell plasticity in resistance to targeted therapy. Nat Rev Drug Discov 19, 39–56. 10.1038/s41573-019-0044-1.

50. Duy, C., Li, M., Teater, M., Meydan, C., Garrett-Bakelman, F.E., Lee, T.C., Chin, C.R., Durmaz, C., Kawabata, K.C., Dhimolea, E., et al. (2021). Chemotherapy Induces Senescence-Like Resilient Cells Capable of Initiating AML Recurrence. Cancer Discov 11, 1542–1561. 10.1158/2159-8290.CD-20-1375.

51. Aggarwal, R., Huang, J., Alumkal, J.J., Zhang, L., Feng, F.Y., Thomas, G.V., Weinstein, A.S., Friedl, V., Zhang, C., Witte, O.N., et al. (2018). Clinical and Genomic Characterization of Treatment-Emergent Small-Cell Neuroendocrine Prostate Cancer: A Multi-institutional Prospective Study. JCO 36, 2492–2503. 10.1200/JCO.2017.77.6880.

52. Uruga, H., Fujii, T., Nakamura, N., Moriguchi, S., Kishi, K., and Takaya, H. (2020). Squamous cell transformation as a mechanism of acquired resistance to tyrosine kinase inhibitor in EGFR-mutated lung adenocarcinoma: a report of two cases. Respirology Case Reports 8, e00521. 10.1002/rcr2.521.

53. Sequist, L.V., Waltman, B.A., Dias-Santagata, D., Digumarthy, S., Turke, A.B., Fidias, P., Bergethon, K., Shaw, A.T., Gettinger, S., Cosper, A.K., et al. (2011). Genotypic and Histological Evolution of Lung Cancers Acquiring Resistance to EGFR Inhibitors. Science Translational Medicine 3, 75ra26–75ra26. 10.1126/scitranslmed.3002003.

54. Wang, Y.-C., Morrison, G., Gillihan, R., Guo, J., Ward, R.M., Fu, X., Botero, M.F., Healy, N.A., Hilsenbeck, S.G., Phillips, G.L., et al. (2011). Different mechanisms for resistance to trastuzumab versus lapatinib in HER2-positive breast cancers--role of estrogen receptor and HER2 reactivation. Breast Cancer Res 13, R121. 10.1186/bcr3067.

55. Luskin, M.R., Murakami, M.A., Keating, J., Winer, E.S., Garcia, J.S., Stahl, M., Wadleigh, M., Flamand, Y., Neuberg, D.S., Galinsky, I., et al. (2023). A Phase I Study of Asciminib (ABL001) in Combination with Dasatinib and Prednisone for BCR-ABL1-Positive ALL and Blast Phase CML in Adults. Blood 142, 965. 10.1182/blood-2023-174246.

56. Keating, A.K., Gossai, N., Phillips, C.L., Maloney, K., Campbell, K., Doan, A., Bhojwani, D., Burke, M.J., and Verneris, M.R. (2019). Reducing minimal residual disease with blinatumomab prior to HCT for pediatric patients with acute lymphoblastic leukemia. Blood Advances 3, 1926–1929. 10.1182/bloodadvances.2018025726.

57. Gökbuget, N., Zugmaier, G., Dombret, H., Stein, A., Bonifacio, M., Graux, C., Faul, C., Brüggemann, M., Taylor, K., Mergen, N., et al. (2020). Curative outcomes following blinatumomab in adults with minimal residual disease B-cell precursor acute lymphoblastic leukemia. Leukemia & Lymphoma 61, 2665–2673. 10.1080/10428194.2020.1780583.

58. Salek, M., Li, N., Chou, H.-P., Saini, K., Jovic, A., Jacobs, K.B., Johnson, C., Lu, V., Lee, E.J., Chang, C., et al. (2023). COSMOS: a platform for real-time morphology-based, label-free cell sorting using deep learning. Commun Biol 6, 971. 10.1038/s42003-023-05325-9.

59. Chan, J.M., Zaidi, S., Love, J.R., Zhao, J.L., Setty, M., Wadosky, K.M., Gopalan, A., Choo, Z.-N., Persad, S., Choi, J., et al. (2022). Lineage plasticity in prostate cancer depends on JAK/STAT inflammatory signaling. Science 377, 1180–1191. 10.1126/science.abn0478.

60. Raghavan, S., Winter, P.S., Navia, A.W., Williams, H.L., DenAdel, A., Lowder, K.E., Galvez-Reyes, J., Kalekar, R.L., Mulugeta, N., Kapner, K.S., et al. (2021). Microenvironment drives cell state, plasticity, and drug response in pancreatic cancer. Cell 184, 6119–6137.e26. 10.1016/j.cell.2021.11.017.

61. Rambow, F., Rogiers, A., Marin-Bejar, O., Aibar, S., Femel, J., Dewaele, M., Karras, P., Brown, D., Chang, Y.H., Debiec-Rychter, M., et al. (2018). Toward Minimal Residual Disease-Directed Therapy in Melanoma. Cell 174, 843–855.e19. 10.1016/j.cell.2018.06.025.

62. Hahn, W.C., Bader, J.S., Braun, T.P., Califano, A., Clemons, P.A., Druker, B.J., Ewald, A.J., Fu, H., Jagu, S., Kemp, C.J., et al. (2021). An expanded universe of cancer targets. Cell 184, 1142–1155. 10.1016/j.cell.2021.02.020.

63. Odejide, O., Weigert, O., Lane, A.A., Toscano, D., Lunning, M.A., Kopp, N., Kim, S., van Bodegom, D., Bolla, S., Schatz, J.H., et al. (2014). A targeted mutational landscape of angioimmunoblastic T-cell lymphoma. Blood 123, 1293–1296. 10.1182/blood-2013-10-531509.

64. Kluk, M.J., Lindsley, R.C., Aster, J.C., Lindeman, N.I., Szeto, D., Hall, D., and Kuo, F.C. (2016). Validation and Implementation of a Custom Next-Generation Sequencing Clinical Assay for Hematologic Malignancies. The Journal of Molecular Diagnostics 18, 507–515. 10.1016/j.jmoldx.2016.02.003.

65. Stockslager, M.A., Malinowski, S., Touat, M., Yoon, J.C., Geduldig, J., Mirza, M., Kim, A.S., Wen, P.Y., Chow, K.-H., Ligon, K.L., et al. (2021). Functional drug susceptibility testing using single-cell mass predicts treatment outcome in patient-derived cancer neurosphere models. Cell Reports 37, 109788. 10.1016/j.celrep.2021.109788.

66. Trombetta, J.J., Gennert, D., Lu, D., Satija, R., Shalek, A.K., and Regev, A. (2014). Preparation of Single-Cell RNA-Seq Libraries for Next Generation Sequencing. Current Protocols in Molecular Biology 107, 4.22.1–4.22.17. 10.1002/0471142727.mb0422s107.

67. Therneau, T. (2024). A Package for Survival Analysis in R.

68. Li, H., and Durbin, R. (2009). Fast and accurate short read alignment with Burrows–Wheeler transform. Bioinformatics 25, 1754–1760. 10.1093/bioinformatics/btp324.

69. McKenna, A., Hanna, M., Banks, E., Sivachenko, A., Cibulskis, K., Kernytsky, A., Garimella, K., Altshuler, D., Gabriel, S., Daly, M., et al. (2010). The Genome Analysis Toolkit: A MapReduce framework for analyzing next-generation DNA sequencing data. Genome Res. 20, 1297–1303. 10.1101/gr.107524.110.

70. DePristo, M.A., Banks, E., Poplin, R., Garimella, K.V., Maguire, J.R., Hartl, C., Philippakis, A.A., del Angel, G., Rivas, M.A., Hanna, M., et al. (2011). A framework for variation discovery and genotyping using next-generation DNA sequencing data. Nat Genet 43, 491–498. 10.1038/ng.806.

71. Cibulskis, K., Lawrence, M.S., Carter, S.L., Sivachenko, A., Jaffe, D., Sougnez, C., Gabriel, S., Meyerson, M., Lander, E.S., and Getz, G. (2013). Sensitive detection of somatic point mutations in impure and heterogeneous cancer samples. Nat Biotechnol 31, 213–219. 10.1038/nbt.2514.

72. McLaren, W., Pritchard, B., Rios, D., Chen, Y., Flicek, P., and Cunningham, F. (2010). Deriving the consequences of genomic variants with the Ensembl API and SNP Effect Predictor. Bioinformatics 26, 2069–2070. 10.1093/bioinformatics/btq330.

73. Bi, W.L., Greenwald, N.F., Ramkissoon, S.H., Abedalthagafi, M., Coy, S.M., Ligon, K.L., Mei, Y., MacConaill, L., Ducar, M., Min, L., et al. (2017). Clinical Identification of Oncogenic Drivers and Copy-Number Alterations in Pituitary Tumors. Endocrinology 158, 2284–2291. 10.1210/en.2016-1967.

74. Li, B., Gould, J., Yang, Y., Sarkizova, S., Tabaka, M., Ashenberg, O., Rosen, Y., Slyper, M., Kowalczyk, M.S., Villani, A.-C., et al. (2020). Cumulus provides cloud-based data analysis for large-scale single-cell and single-nucleus RNA-seq. Nat Methods 17, 793–798. 10.1038/s41592-020-0905-x.

75. Butler, A., Hoffman, P., Smibert, P., Papalexi, E., and Satija, R. (2018). Integrating single-cell transcriptomic data across different conditions, technologies, and species. Nat Biotechnol 36, 411–420. 10.1038/nbt.4096.

76. Weinreb, C., Wolock, S., and Klein, A.M. (2018). SPRING: a kinetic interface for visualizing high dimensional single-cell expression data. Bioinformatics 34, 1246–1248. 10.1093/bioinformatics/btx792.

77. Kotliar, D., Veres, A., Nagy, M.A., Tabrizi, S., Hodis, E., Melton, D.A., and Sabeti, P.C. (2019). Identifying gene expression programs of cell-type identity and cellular activity with single-cell RNA-Seq. eLife 8, e43803. 10.7554/eLife.43803.

78. Gavish, A., Tyler, M., Greenwald, A.C., Hoefflin, R., Simkin, D., Tschernichovsky, R., Galili Darnell, N., Somech, E., Barbolin, C., Antman, T., et al. (2023). Hallmarks of transcriptional intratumour heterogeneity across a thousand tumours. Nature 618, 598–606. 10.1038/s41586-023-06130-4.

79. Kazer, S.W., Aicher, T.P., Muema, D.M., Carroll, S.L., Ordovas-Montanes, J., Miao, V.N., Tu, A.A., Ziegler, C.G.K., Nyquist, S.K., Wong, E.B., et al. (2020). Integrated single-cell analysis of multicellular immune dynamics during hyperacute HIV-1 infection. Nat Med 26, 511–518. 10.1038/s41591-020-0799-2.

80. Kuhn, M. (2008). Building Predictive Models in R Using the caret Package. Journal of Statistical Software 28, 1–26. 10.18637/jss.v028.i05.

81. Wright, M.N., and Ziegler, A. (2017). ranger: A Fast Implementation of Random Forests for High Dimensional Data in C++ and R. Journal of Statistical Software 77, 1–17. 10.18637/jss.v077.i01.

82. Janitza, S., Celik, E., and Boulesteix, A.-L. (2018). A computationally fast variable importance test for random forests for high-dimensional data. Advances in Data Analysis and Classification 12, 885–915.

83. Meyer, P.E. (2008). Information-Theoretic Variable Selection and Network Inference from Microarray Data. In.

84. Badia-i-Mompel, P., Vélez Santiago, J., Braunger, J., Geiss, C., Dimitrov, D., Müller-Dott, S., Taus, P., Dugourd, A., Holland, C.H., Ramirez Flores, R.O., et al. (2022). decoupleR: ensemble of computational methods to infer biological activities from omics data. Bioinformatics Advances 2, vbac016. 10.1093/bioadv/vbac016.

85. Dobin, A., Davis, C.A., Schlesinger, F., Drenkow, J., Zaleski, C., Jha, S., Batut, P., Chaisson, M., and Gingeras, T.R. (2013). STAR: ultrafast universal RNA-seq aligner. Bioinformatics 29, 15–21. 10.1093/bioinformatics/bts635.

86. Danecek, P., Bonfield, J.K., Liddle, J., Marshall, J., Ohan, V., Pollard, M.O., Whitwham, A., Keane, T., McCarthy, S.A., Davies, R.M., et al. (2021). Twelve years of SAMtools and BCFtools. GigaScience 10, giab008. 10.1093/gigascience/giab008.

87. Li, H., Handsaker, B., Wysoker, A., Fennell, T., Ruan, J., Homer, N., Marth, G., Abecasis, G., Durbin, R., and 1000 Genome Project Data Processing Subgroup (2009). The Sequence Alignment/Map format and SAMtools. Bioinformatics 25, 2078–2079. 10.1093/bioinformatics/btp352.

88. Patel, A.P., Tirosh, I., Trombetta, J.J., Shalek, A.K., Gillespie, S.M., Wakimoto, H., Cahill, D.P., Nahed, B.V., Curry, W.T., Martuza, R.L., et al. (2014). Single-cell RNA-seq highlights intratumoral heterogeneity in primary glioblastoma. Science 344, 1396–1401. 10.1126/science.1254257.

89. Tirosh, I., Izar, B., Prakadan, S.M., Wadsworth, M.H., Treacy, D., Trombetta, J.J., Rotem, A., Rodman, C., Lian, C., Murphy, G., et al. (2016). Dissecting the multicellular ecosystem of metastatic melanoma by single-cell RNA-seq. Science 352, 189–196. 10.1126/science.aad0501.

90. Graham, S.M., Vass, J.K., Holyoake, T.L., and Graham, G.J. (2007). Transcriptional Analysis of Quiescent and Proliferating CD34+ Human Hemopoietic Cells from Normal and Chronic Myeloid Leukemia Sources. Stem Cells 25, 3111–3120. 10.1634/stemcells.2007-0250.

