## Supplementary material for "Mutation and cell state compatibility is required and targetable in Ph*+* acute lymphoblastic leukemia minimal residual disease": Table S1

| Characteristic | n | % |
|---|---|---|
| **Age** | | |
| Pediatric | 6 | 40 |
| 18-40 | 4 | 26.7 |
| 41-65 | 3 | 20 |
| 66-80 | 2 | 13.3 |
| **Sex** | | |
| Female | 6 | 40 |
| Male | 9 | 60 |
| **Race** | | |
| White/Non-Hispanic | 8 | 50 |
| White/Hispanic | 1 | 6.3 |
| Black | 3 | 18.8 |
| Asian | 1 | 6.3 |
| Not recorded | 3 | 18.8 |
| **Presenting WBC Count (x10$^3$/µL)** | | |
| >100,000 | 2 | 13.3 |
| 50,001-100,000 | 6 | 40 |
| 10,000-50,000 | 4 | 26.7 |
| <10,000 | 3 | 20 |
| **BCR-ABL Isoform** | | |
| p190 | 10 | 66.7 |
| p210 | 5 | 33.3 |
| **Treatment Phase** | | |
| Diagnosis | 7 | 46.7 |
| Relapse | 7 | 43.8 |
| Refractory | 1 | 6.3 |
| **Prior Therapy** | | |
| Imatinib | 6 | 40 |
| Nilotinib | 2 | 13.3 |
| Dasatinib | 4 | 26.7 |
| Ponatinib | 4 | 26.7 |
| Any TKI | 7 | 46.7 |
| Cytotoxic chemotherapy | 6 | 40 |
| Allogeneic HSCT | 3 | 20 |
| Autologous HSCT | 1 | 6.7 |
| **Patient ABL1 Mutation** | | |
| Any | 3 | 20 |
| T315I | 1 | 6.7 |
| None | 3 | 20 |
| Sequencing not done | 9 | 60 |
| **IKZF1 deletion** | | |
| Yes | 9 | 60 |
| No | 6 | 40 |
| **Deletion 7p** | | |
| Yes | 0 | 0 |
| No | 13 | 86.7 |
| Not determined | 2 | 13.3 |
| **Hyperdiploid** | | |
| Yes | 3 | 20 |
| No | 10 | 66.7 |
| Not determined | 2 | 13.3 |
| **Trisomy 21** | | |
| Yes | 4 | 26.7 |
| No | 9 | 60 |
| Not determined | 2 | 13.3 |

**Table 1. Clinical characteristics of patients** whose tumors were used to generate PDX models.
