## Supplementary material for "Mutation and cell state compatibility is required and targetable in Ph*+* acute lymphoblastic leukemia minimal residual disease": Table S2

| PDX Name | Patient Age | Patient Sex | Patient Race/Ethnicity | Patient Initial WBC Count ($x10^3/\mu L$) | Patient Phase of Disease at Time of Xenografting | Patient Prior Therapy | Tumor Karyotype | BCR-ABL1 Isoform | Patient Tumor Immunophenotype Positive | Patient Tumor Immunophenotype Negative | Patient Tumor Mutations | Patient Tissue Xenografted | PDX Mutations | PDX IKZF1 Deletion | Patient Tumor Deletion 7p | Patient Tumor Deletion 9p | Patient Tumor Hyperdiploid | Patient Tumor Trisomy 21 |
|---|---|---|---|---|---|---|---|---|---|---|---|---|---|---|---|---|---|---|
| DFAB-92612 | 28 | M | White | 56000 | Refractory | Induction [Prednisone, doxorubicin, vincristine, HD-MTX, L-asparaginase, IT-cytarabine] | 46,XY,t(9;22)(q34;q11.2)[11]/46,XY[9] | p210 | CD34, CD45 (dim), HLA-DR, TdT, CD13, CD33, CD10, CD19 | CD20, surface immunoglobulin, kappa and lambda light chains, CD117, T lymphoid, and monocytic markers | Not sequenced | Bone Marrow | KRAS Y64C | Yes | No | No | No | No |
| CBAB-30198 | 11.7 | M | White | 132800 | Relapse | Induction [details N/A]<br>Consolidation [cyclophosphamide, cytarabine, 6-MP, IT-MTX], dasatinib | 45,XY,add(9)(q34),psudic(19;9)(p13.3;p13),<br>der(22)t(9;22)(q34;q11.2)[1]/<br>45,sl,t(7;19)(q11.2;q13.3)[4]/<br>45,sl,t(X;7)(q21.1;p22)[3]/46,XY[12] | p210 | CD45 (dim), CD19, CD10, CD34, HLA-DR, CD9, CD58, CD44 and partial CD20, CD22, CD81, CD73, TdT | CD2, CD7, CD15, CD38, CD117, CRLF2, surface immunoglobulin light chains, CD13/CD33 mostly negative | None | Peripheral Blood | None | Yes | No | No | No | No |
| DFAB-96061 | 62 | F | White | 73690 | Relapse | Induction [hyper-CVAD], imatinib<br>dasatinib for MRD<br>Consolidation [allogeneic HSCT],<br>post-transplant dasatinib<br>Salvage [nilotinib] \| Consolidation [DLI] | 46,XX,t(9;22)(q34;q11.2,del(11)([11.2))[1]/<br>46,XY[19] | p210 | CD45 (dim), CD34, HLA-DR, TdT, CD19, CD10, CD22 (dim), CD33 (dim), CD13 (dim subset) | CD20, surface immunoglobulin, T cell and other myeloid markers | ABL1 Y253H 16% of 167 reads<br>ABL1 F311L 7% of 234 reads<br>ABL1 E255V VAF not specified<br>SETD2 E1412* 11% of 225 reads<br>ROS1 T804N 20% of 185 reads | Leukapheresis | ABL1 Y253H<br>ABL1 F311L<br>ABL1 T315I | Yes | No | No | No | No |
| DFAB-25157 | 62 | F | White | 73690 | Relapse | Induction [hyper-CVAD], imatinib<br>dasatinib for MRD<br>Consolidation [allogeneic HSCT],<br>post-transplant dasatinib<br>Salvage [nilotinib] \| Consolidation [DLI]<br>Salvage [vincristine x 1] | 46,XX,t(9;22)(q34;q11.2,del(11)([11.2))[1]/<br>46,XY[19] | p210 | CD45 (dim), CD34, HLA-DR, TdT, CD19, CD10, CD22 (dim), CD33 (dim), CD13 (dim subset) | CD20, surface immunoglobulin, T cell and other myeloid markers | ABL1 Y253H 16% of 167 reads<br>ABL1 F311L 7% of 234 reads<br>ABL1 E255V (VAF not specified)<br>SETD2 E1412* 11% of 225 reads<br>ROS1 T804N 20% of 185 reads | Bone Marrow | ABL1 Y253H 52.6% of 133 reads<br>MKI67 Q2084P 50.0% of 326 reads | Yes | No | No | No | No |
| DFAB-13601 | 59 | F | Unavailable | 27700 | Relapse | Induction [cyclophosphamide, daunorubicin, vincristine, prednisone, L-asparaginase]<br>Early Intensification [IT-MTX, cyclophosphamide, 6-MP, cytarabine, vincristine, L-asparaginase], imatinib<br>Consolidation [allogeneic HSCT] | 47,XX,+2,i(8)(q10),der(9)t(9;22)(q34;q11.2),<br>del(16)(q22),idicder(22)t(9;22)(q34;q11.2)[13]/<br>47,XX,+2,i(8)(q10),t(9;22(q34;q11.2),<br>del(16)(q22)[1]/<br>46, XX[6] | p190 | CD45(DIM), HLA-DR, CD34, TDT, B lymphoid markers CD19, CD20, CD10 | Surface immunoglobulin,CD3, CD5, CD7, CD13, CD33, CD117, CD15 | Not sequenced | Bone Marrow | ABL1 T315I 47.4% 230 reads<br>PRKDC V1479L 52.9% of 221 reads<br>PRKDC L1707Q 50.9% of 226 reads<br>KMT2C Y987H 13.0% of 600 reads<br>KMT2C S990G 6.1% of 588 reads<br>KMT2C 5.5% 860 reads | Yes | No | No | No | No |
| SFAB-62876 | 36 | M | Unavailable | 99000 | Relapse | Induction [cyclophosphamide, daunorubicin, vincristine, prednisone, L-asparaginase]<br>Consolidation [hyper-CVAD 1B]<br>Re-induction [hyper-CVAD "A", rituximab]<br>Re-induction [HAM]<br>Salvage [FLAG-Ida, imatinib] | 45~46,XY,del(9)(p?21)t(9;22)(q34;q11.2)[cp2]/<br>45~46,sl,add(11)(q23),-19, +mar[cp14]/<br>45~46,sdl1,add(10)(p1?3)[cp2]/46,XY[2] | p210 | Missing | Missing | GNB1 K89E (VAF not specified) | Bone Marrow | GNB1 K89E 52.7% of 184 reads | Yes | No | Yes | No | No |
| CBAB-75728 | 5 | M | Black | 32000 | Relapse | Induction [prednisone, doxorubicin, vincristine, HD-MTX, PEG-asparaginase, IT-MAH, IT-MTX], imatinib | Full report not available | p190 | Missing | Missing | Not sequenced | Peripheral Blood | PTPN11 M504V | Yes | ND | Yes | ND | ND |
| HCAB-89433 | 51 | F | Black | 3000 | Relapse | Induction [hyper-CVAD + imatinib]<br>Consolidation [autologous HSCT]<br>Salvage [ponatinib] | Full report not available | p190 | CD123, CD303, BDCA4 (CD304/neutropilin1), CD56, CD4, CD45low DR, CD34, CD7, CD38, | BDCA1-(CD1c), CD33, CD56 | None | Bone Marrow | KRAS G13D | Yes | ND | ND | ND | ND |
| DFAB-54880 | 31 | F | Black | 9900 | Untreated | None | 46,XX,t(9;22)(q34;q11.2)[1]/45,XX,idem,<br>del(3)(p11), der(9;12)(q10;q10)[7]/<br>45,XX, idem,del(5)(q13),der(9;12),<br>der(11)t(5;11)(q13;q21)[7]/<br>46,XX[5] | p190 | CD45 (dim), CD34, HLA-DR, TDT, CD19, CD10 | CD33, CD20, surface immunoglobulin, CD3, CD5, CD7, CD13, CD117, CD15, CD56, CD11b, CD14 and CD64 | None | Bone Marrow | None | Yes | No | No | No | No |
| CBAB-12402 | 2.8 | M | Asian | 32600 | Untreated | None | 48XY,+?18,+21 | p190 | CD45 (dim), TdT, CD34, CD38 (variable, dim), CD58, HLA-DR, CD19, CD20 (variable), surface CD22 (dim), CD10 (bright) and CD9 | Surface immunoglobulin light chains, surface IgM as well as other myelomonocytic (CD13, CD33, CD117) and T-cell (CD2, CD7, CD56) markers | Unknown | Peripheral Blood | None | No | No | No | No | Yes |
| DFAB-74952 | 44 | F | White | 54000 | Untreated | None | 42-46,XX,-8,i(8)(q10)[6],t(9;22)(q34;q11.2),<br>der(16)t(8;16)(q12;q12),+der(16)t(8;16)<br>[6],+der(22)t(9;22)[2][cp16]/46,XX[4] | p190 | CD45+,CD34+,HLA-DR+,TdT,CD19,CD10,CD22 | CD20,surface immunoglobulin,T-cell markers,myeloperoxidase | Not sequenced | Bone Marrow | None | ND | No | ND | No | No |
| DFAB-90281 | 59 | M | Unavailable | 69500 | Untreated | None | 46,XY,t(9;22)(q34;q11.2)[9]/45,XY,idem,-7[7] | Missing | CD34,CD45,HLA-DR,CD13(dim),CD33(dim),CD10,CD19, TdT | CD20,surface immunoglobulin, T lymphoid and monocytic markers | Not sequenced | Bone Marrow | Not sequenced | ND | No | ND | No | No |
| CBAB-99093 | 14.2 | M | White | 25800 | Untreated | None | Failed; BCR-ABL1 by PCR; del(9p) by FISH | p190 | Missing | Missing | Not sequenced | Bone Marrow | PTPN11 M504V | No | ND | Yes | ND | ND |
| CBAB-75914 | 16.1 | M | Hispanic | 70100 | Untreated | None | 56XY,+X,+2,+4,+5,+6,t(9;22)(q34;q11.2),+11,+14,<br>+14,+21.der(22)t(9;22) | p190 | CD45 (dim), TdT, CD34 (variable), CD38 (dim variable), CD58, HLA-DR, CD19, CD20 (heterogeneous), sCD22 (dim), CD10 and CD9 (variable) | Surface immunoglobulin light chains as well as surface IgM and other myelomonocytic (CD13, CD15, CD33, CD117) and T-cell (CD2, CD7, CD56) markers | Unknown | Peripheral Blood | None | No | No | No | Yes | Yes |
| DFAB-62208 | 40 | M | White | 6800 | Untreated | None | 59,XY,+X,+2,+4,+5,+6,add(6)(q2?5),+8,<br>t(9;22)(q34;q11.2),+10,+11,+14,+17,+17,+18,+21<br>, +der(22)t(9;22),+mar[cp20] | p190 | CD34, CD45, HLA-DR, CD19,CD10, TdT | CD20, surface immunoglobulin, T-lymphoid, myeloid, monocytic markers | Not sequenced | Bone Marrow | None | No | No | No | Yes | Yes |
| CBAB-72204 | 2.3 | M | White | 175100 | Untreated | None | 55XY,+X,+4,+6,t(9;22)(q34;q11.2),<br>+13,+14,+14,+17,+21,+der(22)t(9;22) | p190 | Missing | Missing | Unknown | Bone Marrow | None | No | No | No | Yes | Yes |

**Table 2. Characteristics of PDX models** of BCR-ABL ALL and patients whose tumors were used to generate them. ND = not determined.
